# *NITROGEN LIMITATION ADAPTATION* functions as a negative regulator of Arabidopsis immunity

**DOI:** 10.1101/2021.12.09.471910

**Authors:** Beatriz Val-Torregrosa, Mireia Bundó, Tzyy-Jen Chiou, Victor Flors, Blanca San Segundo

**Affiliations:** Centre for Research in Agricultural Genomics (CRAG) CSIC-IRTA-UAB-UB. Edifici CRAG, Carrer de la Vall Moronta. Campus Universitat Autònoma de Barcelona (UAB), Bellaterra (Cerdanyola del Vallés), 08193 Barcelona, Spain; Agricultural Biotechnology Research Center, No. 128, Sec. 2, Academia Road, Nankang, Taipei 115, Taiwan; Departamento de Ciencias Agrarias y del Medio Natural, Escuela Superior de Tecnología y Ciencias Experimentales, Universitat Jaume I, Spain; Consejo Superior de Investigaciones Científicas (CSIC), Barcelona, Spain

**Keywords:** Arabidopsis, callose, camalexin, Jasmonic Acid, Immnunity, miR827, Phosphate, *Plectosphaerella cucumerina*, Salicylic Acid

## Abstract

**Background:** Phosphorus is an important macronutrient required for plant growth and development. It is absorbed through the roots in the form of inorganic phosphate (Pi). To cope with Pi limitation, plants have evolved an array of adaptive mechanisms to facilitate Pi acquisition and protect them from stress caused by Pi starvation. The NITROGEN LIMITATION ADAPTION (NLA) gene plays a key role in the regulation of phosphate starvation responses (PSR), its expression being regulated by the microRNA miR827. Stress caused by Pi limiting conditions might also affect the plant’s response to pathogen infection. However, cross-talk between phosphate signaling pathways and immune responses remains unclear.

**Results:** In this study, we investigated whether *NLA* plays a role in Arabidopsis immunity. We show that loss-of-function of *NLA* and *MIR827* overexpression causes an increase in phosphate (Pi) content which results in resistance to infection by the fungal pathogen *Plectosphaerella cucumerina*. The *nla* mutant plants accumulated callose in their leaves, a response that is also observed in wild-type plants that have been treated with high Pi. We also show that pathogen infection and treatment with fungal elicitors is accompanied by transcriptional activation of *MIR827* and down-regulation of *NLA*. Upon pathogen challenge, *nla* plants exhibited higher levels of the phytoalexin camalexin compared to wild type plants. Camalexin level also increases in wild type plants treated with high Pi. Furthermore, the *nla* mutant plants accumulated salicylic acid (SA) and jasmonic acid (JA) in the absence of pathogen infection whose levels further increased upon pathogen.

**Conclusions:** This study shows that *NLA* acts as a negative regulator of Arabidopsis immunity. Overaccumulation of Pi in *nla* plants positively affects resistance to infection by fungal pathogens. This piece of information reinforces the idea of signaling convergence between Pi and immune responses for the regulation of disease resistance in Arabidopsis.

## Background

Phosphorous (P) is one of the most important macronutrients required for plant growth and development. It is a key component of fundamental biomolecules, such as nucleic acids phospholipids, and ATP, the principal molecule for storing and transferring energy in cells. P is also a central regulator in numerous metabolic reactions that also functions in signaling pathways and modulation of protein activity. Plants absorb phosphorus from the soil in the form of inorganic form (Pi). Although P levels in soils are usually high, its bioavailability is often extremely low which represents a limiting factor for plant growth.

To cope with Pi limitation, plants have evolved multiple molecular mechanisms that allows the plant to enhance the acquisition of external Pi, collectively known as the Pi starvation response (PSR) (Yang and Finnegan, 2010; Puga *et al*., 2017; Chien et al., 2018; Wang *et al*., 2021). In Arabidopsis, the transcription factor PHR1 (PHOSPHATE STARVATION RESPONSE 1) is a central regulator of Pi starvation responses leading to the induction of phosphate transporter genes of the *PHT1* family (Bustos *et al*., 2010; Nilsson *et al*., 2012; Guo *et al*., 2015). *AtPHR1* regulates a number of Pi starvation-induced genes through binding a P1BS (PHR1 specific binding sequence) *cis*-element, which is present in the promoter regions of Pi starvation-induced genes (Rubio *et al*., 2001). PHR1 also induces the expression of two microRNAs (miRNAs), miR399 and miR827, which negatively regulate the expression of *PHO2* (*PHOSPHATE 2*, a ubiquitin E2 conjugase) and *NLA* (*NITROGEN LIMITATION ADAPTATION*, a ubiquitin E3 ligase), respectively (Delhaize and Randall, 1995; Lin *et al*., 2013). PHO2 and NLA mediate the ubiquitination and degradation of the plasma-membrane-localized PHT1 transporters. In this way, miRNA-mediated down-regulation of *PHO2* and *NLA* relieves the negative regulation over PHT1 transporters, thus, increasing Pi uptake in the roots (Bari *et al*., 2006; Kant *et al*., 2011). Whereas considerable progress has been made in characterizing the regulatory mechanisms that underlie adaptation of plants to Pi limiting conditions, less is known about adaptive mechanisms to Pi excess condition.

On the other hand, an adequate supply of Pi is also important in disease resistance. Evidence support that PSR can interact with the plant immune system in Arabidopsis (Castrillo *et al*., 2017; Chan *et al*., 2021). Under low Pi conditions, the *PHR1* transcription factor target genes involved in the JA- and/or SA-dependent pathways for suppression of immune responses (Castrillo et al., 2017). As a consequence, *phr1* Arabidopsis mutants exhibit resistance to infection by bacterial and oomycete pathogens (*P. syringae* DC3000 and *Hyaloperonospora arabidopsidis*, respectively) (Castrillo *et al*., 2017). Also, PHR1 negatively regulates the immune response triggered by the bacterial elicitor peptide flagellin22 (flg22). This study also led the authors to propose that the Arabidopsis plants prioritizes Pi stress responses over defense (Castrillo *et al*., 2017). PHR1, however, does not necessarily comprommises the entire immune system as immune responses are still activated by certain pathogens, even in low Pi conditions (Hacquard et al., 2016). In other studies, transgenic expression of a phytoplasma effector (SAP11) in Arabidopsis was found to trigger Pi starvation responses that are mainly dependent of *PHR1* (Lu *et al*., 2014). The SAP11 transgenic plants overaccumulated Pi in leaves and were more susceptible to *P*.*syringae* pv.tomato DC3000 infection (Lu *et al*., 2014). In other studies, high Pi fertilization and miR399 overexpression, and subsequent Pi accumulation, was found to compromise expression of the defense-related genes OsPR1 and OsPBZ1 in rice, thus, resulting in a phenotype of susceptibility to infection by the blast fungus (Campos-Soriano *et al*., 2020). This piece of information reinforces the idea of cross-talk between Pi signaling and immune signaling in plants. However, the molecular mechanisms by which Pi excess might regulate plant immunity still remain unclear.

The plant immune system consists of interconnected processes that are induced upon recognition of the pathogen. Depending on the molecules that are recognized by the host plant, two immune systems have been defined. Plants recognize pathogen epitopes, known as Pathogen-Associated Molecular Patterns (PAMPs, also known as elicitors). This recognition activates a general defense response referred to as PAMP-triggered immunity (PTI) leading to the induction of defense-related genes, such as *Pathogenesis-Related* (*PR*) genes (Jones and Dangl, 2006; Boller and Felix, 2009; Li *et al*., 2020). Callose deposition is a hallmark of the induction of PTI presumably to fortify the plant cell wall at pathogen infection sites. Pathogens adapted to their host have evolved effectors that are delivered to plant cells and suppress PTI leading to disease susceptibility. During co-evolution, plants have evolved another immune system in which pathogen effectors, or host proteins modified by these effectors, are recognized by proteins encoded by resistance (R) genes, called Effector-triggered Immunity (ETI). Whereas PTI contributes to resistance to diverse pathogens, ETI is pathogen strain or race specific. Phytohormones also play an essential role in the regulation of plant immune responses. The involvement of salicylic acid (SA), jasmonic acid (JA), and ethylene (ET) in the regulation of defense responses to pathogens in plants has long been recognized (Denancé *et al*., 2013; Berens *et al*., 2017; Aerts *et al*., 2021). Synergistic and antagonistic interactions between hormone signaling pathways allow fine-tune responses to different pathogens.

Phytoalexins are low molecular weight antimicrobial compounds that take part in the defense system used by plants against pathogens (Ahuja *et al*., 2012). The major phytoalexin involved in resistance against pathogens in Arabidopsis is camalexin, a sulfur-containing tryptophan-derived secondary metabolite. Camalexin accumulation has been correlated with resistance to necrotrophic fungi such as *Plectosphaerella cucumerina, Botrytis cinerea* and *Alternaria brassicicola* (Ferrari *et al*., 2003; Nafisi *et al*., 2007; Sanchez-Vallet *et al*., 2010).

In this work, we investigated whether *NLA* and miR827, which function downstream of *PHR1* in the PSR, play a role in the resistance to infection by fungal pathogens in Arabidopsis. As previously mentioned, miR827 down-regulates *NLA* expression in Pi-starved Arabidopsis plants, the *NLA* gene encoding a RING-type ubiquitin ligase responsible of ubiquitination of PHT1 transporters (Hsieh *et al*., 2009; Kant *et al*., 2011). Loss-of-function of *NLA* results in overaccumulation of PHT1 phosphate transporters, thereby increasing Pi uptake (Kant *et al*., 2011). Even though the *NLA* gene was originally described as a positive regulator in the adaptive response of Arabidopsis plants to nitrogen (N) limitation (Peng *et al*., 2007), it is widely recognized that *NLA* also plays a pivotal role in the regulation of Pi homeostasis (Lin *et al*., 2013).

In this study, we show loss-of-function of *nla* and miR827 overexpressor plants accumulate Pi, these plants also exhibiting resistance to infection by fungal pathogens. Transcriptional activation of *MIR827*, and subsequent down-regulation of *NLA*, occurs in response to pathogen infection and treatment with fungal elicitors. Upon pathogen infection, *nla* plants showed higher deposition of callose compared with wild-type plants. We also show that loss-of-function of *NLA*, as well as treatment with high Pi, is accompanied by accumulation of camalexin in leaves. *nla* plants also exhibited higher levels of SA and JA in the absence of pathogen infection. Overall, the results presented here suggest that *NLA* plays a negative role in Arabidopsis immunity.

## Results

### Resistance to infection by fungal pathogens in *nla* and miR827 overexpressor plants

To investigate whether *NLA* and miR827 play a role in resistance to fungal infection in Arabidopsis, we initially examined disease resistance in *nla* mutant and miR827 overexpressor plants (henceforth miR827 OE). As miR827 down-regulates *NLA* expression, the loss-of-function mutation of *nla* is expected to have a disease phenotype similar to that of transgenic lines overexpressing miR827. The *nla* mutant used for these studies, *nla-1* (Col-0 background), is a RING-domain deletion mutant caused by deletion of the third and fourth exon which was described elsewhere (Peng *et al*., 2007; Kant *et al*., 2011; Lin *et al*., 2013) (**Fig. S1A**). Compared with wild type plants, the *nla* plants accumulated Pi in their leaves and showed down-regulation of *NLA* expression which is consistent with results previously reported (Lin *et al*., 2013) (**Fig. S1B**). As for miR827 OE plants, the production and characterization of these plants was previously described (Kant et al., 2011). As expected, miR827 OE plants accumulated pre-miR827 transcripts and had reduced *NLA* expression, these plants also accumulating Pi in their leaves (**Fig. S1C**).

The *nla* and miR827 OE plants were tested for resistance to infection by the fungus *Plectosphaerella cucumerina*. The Arabidopsis/*P*.*cucumerina* pathosystem is a well-established model for studies on basal resistance to infection by necrotrophic fungi (Sanchez-Vallet *et al*., 2010). Three-week-old plants were inoculated with P. cucumerina spores. At this developmental stage, no phenotypic differences were observed between *nla*, miR827 OE and wild type plants (**Fig. S2**). At a later developmental stage, however, miR827 OE plants displayed a reduced size compared with wild type plants which is in agreement with results reported by Hewezi *et al*. (2016). Accordingly, to avoid that differences in development caused by miR827 OE could interfere with disease resistance assays, all infection experiments were carried out on 3 week-old plants (wild type, miR399, *nla* plants).

Interestingly, *nla* and miR827 OE plants consistently showed enhanced resistance to infection by *P. cucumerina* compared with the wild-type plants (**Fig. 1A**). Quantification of fungal biomass and determination of plant survival confirmed resistance to *P. cucumerina* in both *nla* and miR827 OE plants (**Fig. 1B**). Moreover, trypan blue staining of infected leaves revealed extensive fungal growth in leaves of wild-type plants, but not in leaves of *nla* plants (**Fig. 1C**).

**Fig. 1.**
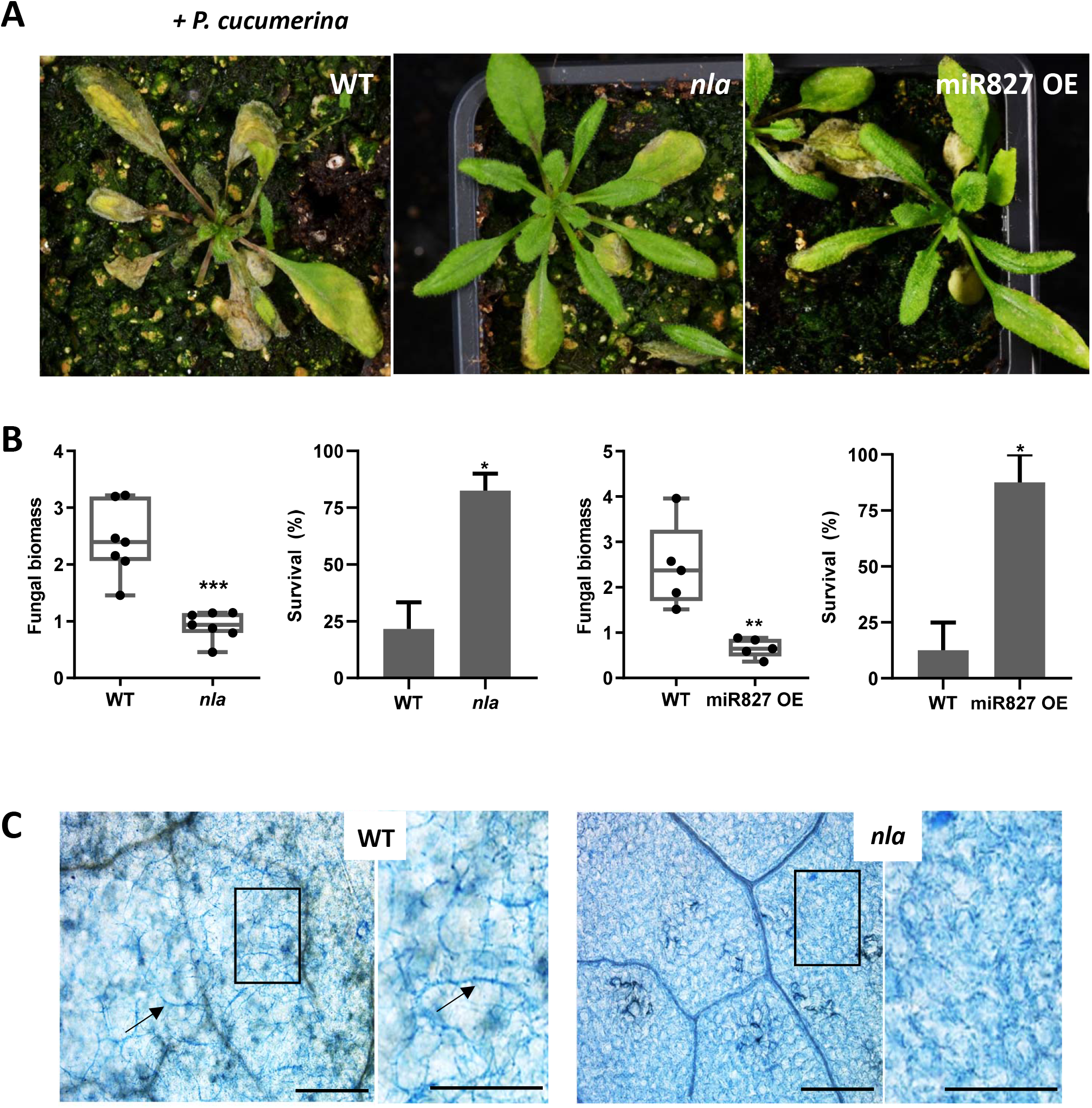
Resistance of *nla* and miR827 OE plants to infection by necrotrophic fungal pathogen *P. cucumerina*. Three-week-old plants were mock-inoculated or inoculated with *P. cucumerina* (5·10^5^ spores/ml). Three independent infection experiments were carried out with similar results (at least 24 plants per genotype each experiment). **A**. Phenotype of *P. cucumerina*-infected wild-type, *nla* and miR827 OE plants. Images were taken at 7 days after inoculation (dpi). **B**. Fungal biomass and survival of *nla* and miR827 OE plants. Quantification of fungal DNA was performed by qPCR using specific primers of *P. cucumerina* (Soto-Suárez *et al*., 2017) at 7 dpi. Survival was determined at 7 dpi. **C**. Trypan blue staining of leaves from *P. cucumerina*-infected *nla* plants at 7 dpi. Arrows indicate fungal hyphae. Scale bars represent 200 µm. Higher magnifications are shown (right panels; bars, 100 µm).

The *nla* plants also exhibited resistance to infection by the hemibiotrophic fungus *Colletotrichum higginsianum*, which was confirmed by the amount of fungal biomass and percentage of diseased leaves (**Fig. S3A, B)**. *C. higginsianum* causes the anthracnose leaf spot disease of *Brassica* species, including *A. thaliana* (O’Connell *et al*., 2004). Subsequent studies in this work were conducted on *nla* mutant plants during interaction with the fungus *P. cucumerina*.

### Callose accumulation in *nla* plants in response to *P. cucumerina* infection

Callose accumulation is known to be important factors in the Arabidopsis defense response to infection by *P. cucumerina* and represents the first line of defense defense that avoid fungal penetration (Luna *et al*., 2011; Pastor et al., 2013: Wang *et al*., 2021). Fungal elicitors are also potent PAMPs for inducing callose deposition.

In this work, aniline blue was used to visualize callose accumulation in leaves of wild-type and *nla* plants that have been inoculated with *P. cucumerina* spores, or mock-inoculated. Neither wild-type nor *nla* plants showed callose deposition in the absence of pathogen infection (mock conditions) (**Fig. 2A**, Mock). Upon challenge with *P. cucumerina*, both wild type and *nla* plants accumulated callose. Notably, *nla* plants infected with *P. cucumerina* showed significantly higher callose deposits compared with *P. cucumerina*-infected wild-type plants (**Fig. 2A**). Quantification of callose deposition confirmed higher accumulation in *nla* plants than in wild-type plants during pathogen infection (**Fig. 2A**, right panel). The observation that callose does not accumulates in *nla* plants in the absence of pathogen infection suggest that this immune response is not constitutively active in *nla* plants. Increased callose accumulation in *nla* plants during pathogen might well contribute to arrest pathogen growth in these plants.

**Fig. 2.**
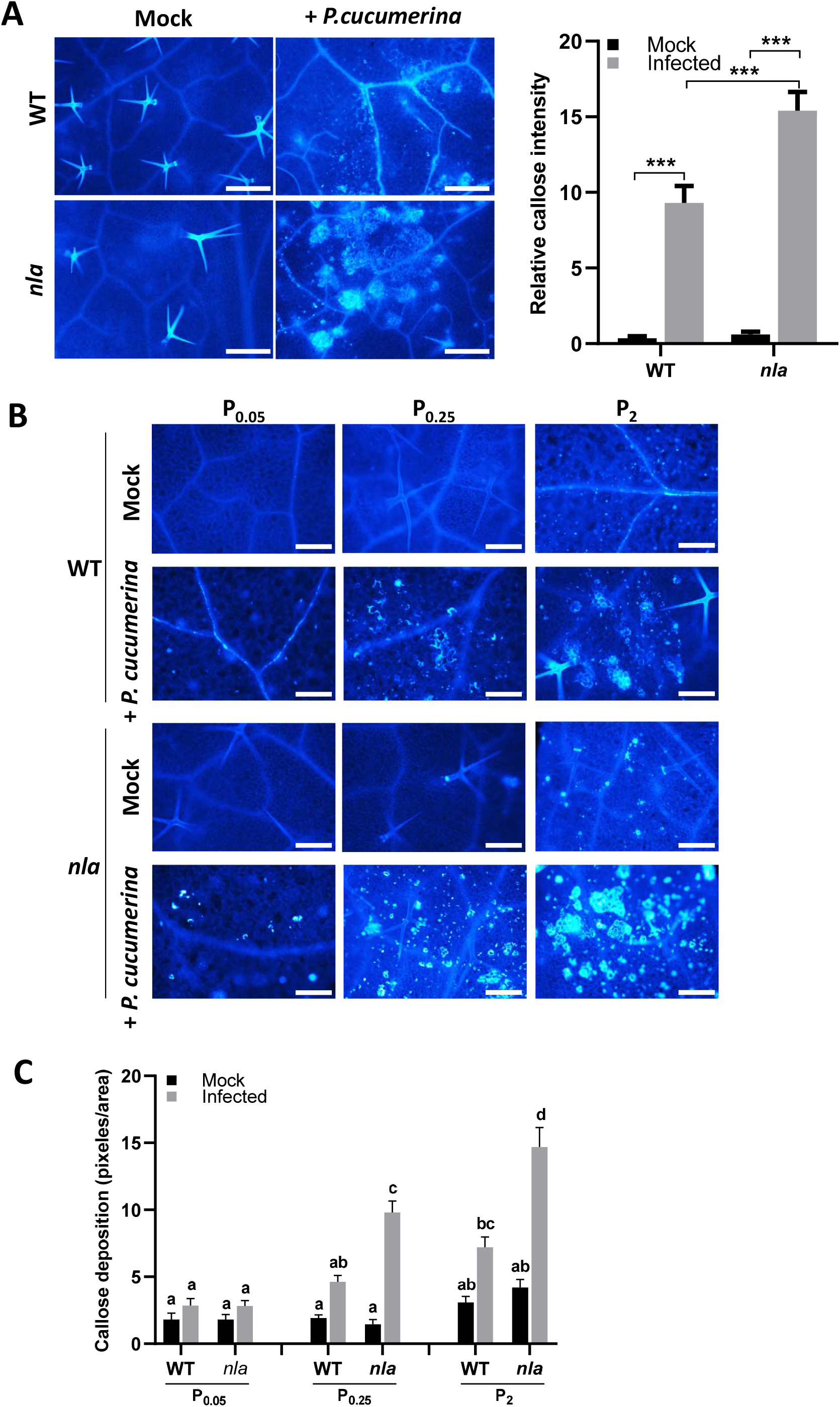
Callose deposition in *P. cucumerina*-infected leaves of wild-type and *nla* plants. Wild-type and *nla* plants were inoculated with *P. cucumerina* spores (5 × 10^5^ spores/ml), or mock-inoculated. Aniline blue staining was carried out at 48 hpi. Representative results from one of three independent infection experiments that gave similar results are shown (at least 12 plants each experiment). **A**. Micrographs of aniline blue stained leaves. (left panel). Bars represent 200 μm. Right panel, pathogen-induced callose deposition was quantified by determining the relative number of fluorescent pixels on digital micrographs of infected leaves (right panel). Bars represent mean ± SEM. **B**. Callose accumulation in Arabidopsis plants that have been grown *in vitro* under increasing Pi supply for 7 days (0.05 mM, 0.25 mM and 2 mM Pi), and then inoculated with *P. cucumerina* spores, or mock-inoculated. **C**. Quantification of callose deposition in wild-type and *nla* plants (same conditions as in B).

Knowing that *nla* plants accumulated more callose after challenge with *P. cucumerina*, these plants also accumulating Pi in leaves, it was of interest to determine whether callose accumulation is affected by Pi supply. Towards this end, wild-type and *nla* plants were grown *in vitro* under different Pi concentrations (0.05 mM, 0.25mM and 2 mM Pi (hereinafter P_0.05_, P_0.25_ and P_2_, respectively). Measurement of Pi content confirmed higher leaf Pi content when increasing Pi supply to plants) in wild type and *nla* plants (**Fig. S4**). Aniline blue staining of non-infected wild type and *nla* plants revealed small fluorescent callose spots at the highest Pi condition (2 mM Pi), which had a larger size in *nla* plants compared with wild-type plants (**Fig. 2B**, mock-inoculated wild type, mock-inoculated *nla* plants). Upon pathogen challenge, callose deposits were more abundant and had a larger size in *P. cucumerina*-infected *nla* plants grown under high Pi supply (P2 plants) (**Fig. 2B**, *P. cucumerina*-inoculated wild type and *P. cucumerina*-inoculated *nla* plants). Quantification of callose deposition confirmed higher accumulation in *nla* plants compared with WT plants during pathogen infection (**Fig. 2C**). Thus, these results demonstrated that fungal infection is accompanied by higher callose deposition in *nla* plants compared with wild type plants.

### *NLA* and *MIR827* expression is regulated during fungal infection

The genome of *Arabidopsis thaliana* contains a single copy of the *MIR827* gene, whose expression is induced in Pi limiting conditions, resulting in down-regulation of *NLA* expression (Hsieh *et al*., 2009). In this work, we investigated whether infection by the fungal pathogen *P. cucumerina* has an effect on *MIR827* and *NLA* expression. As shown in **Fig. 3A**, the accumulation of miR827 precursor and mature transcripts significantly increased in *P. cucumerina*-infected plants compared with non-infected plants, whereas *NLA* was repressed in the fungal-infected plants.

**Fig. 3.**
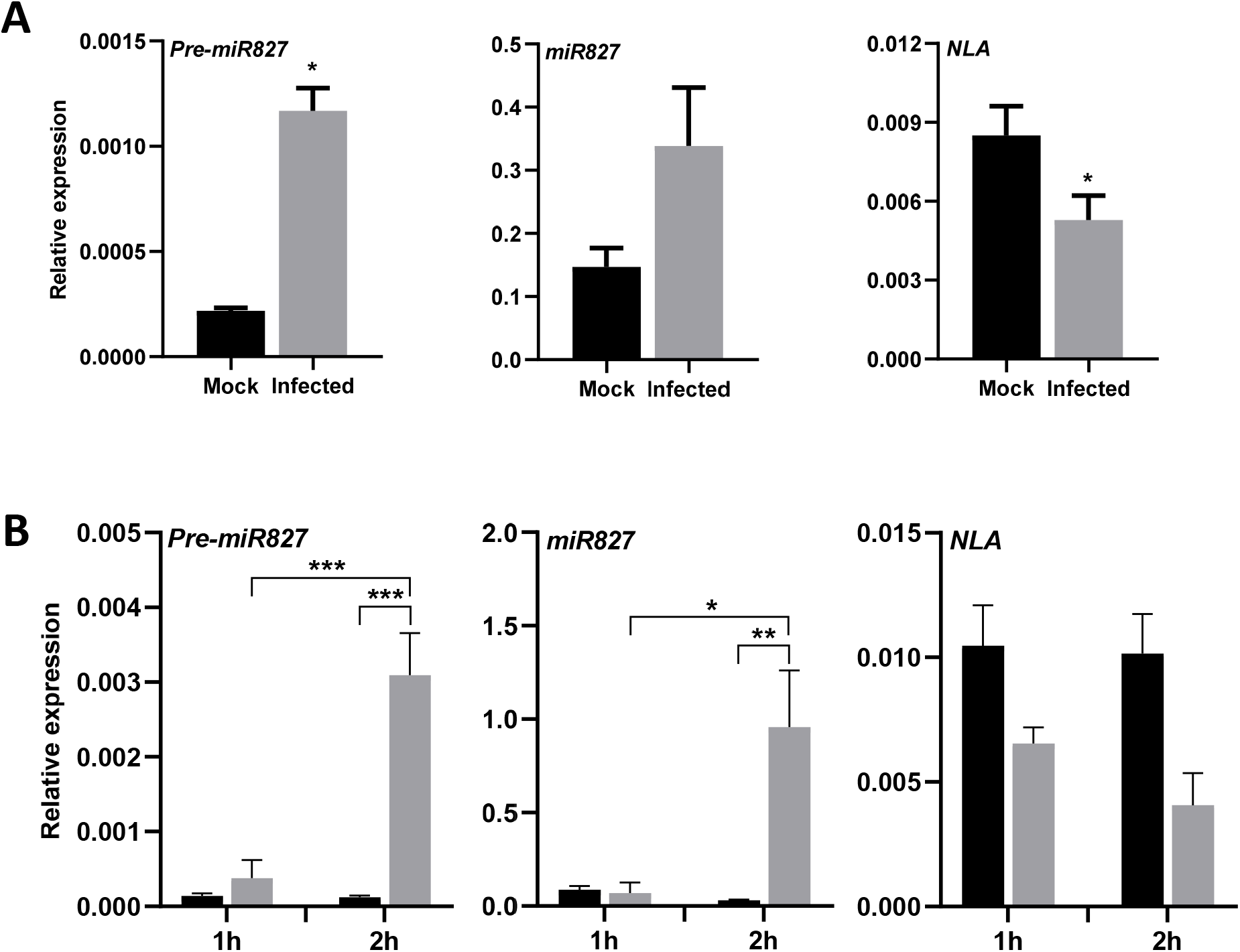
*MIR827* and *NLA* expression in response to infection with *P. cucumerina* and treatment with *P. cucumerina* elicitors. Plants were grown in soil for three weeks. Accumulation of pre-miR827, mature miR827 and *NLA* transcripts in response to inoculation with *P. cucumerina* spores (5·10^5^ spores/ml) at 48hpi (**A**) or treatment with elicitors obtained from this fungus (300 µg/ml elicitors) (**B**). The level of transcripts was determined by RT-qPCR (pre-miR827) and stem-loop RT-qPCR (miR827). Black bars, mock-inoculated plants. Grey bars, *P. cucumerina*-inoculated or elicitor-treated plants. Histograms show the mean ± SEM. Statistical significance was determined by ANOVA (\**p* ≤ 0.05, \*\**p* ≤ 0.01 and \*\*\**p* ≤ 0.001).

On the other hand, it is well known that perception of elicitors (or PAMPs) triggers the induction of PTI. To note, treatment with a crude preparation of *P. cucumerina* elicitors was found to be accompanied by up-regulation of *MIR827* expression and down-regulation of *NLA* expression (**Fig. 3B)**.

From these results it is concluded that *MIR827* expression is up-regulated during fungal infection and treatment with fungal elicitors. Since *NLA* is the target gene of miR827, induction of *MIR827* expression during infection (or elicitor treatment) might account, at least in part, for the reduced expression of *NLA*.

To get further support of elicitor-responsiveness of *MIR827*, we generated transgenic lines expressing the *GUS* reporter gene under the control of the *MIR827* promoter (*MIR827prom::GUS* lines). As control, transgenic Arabidopsis plants expressing the *GUS* reporter gene under the control of the *35S Cauliflower Mosaic Virus 35S* promoter (*35Sprom::GUS* plants) were also examined. Arabidopsis plants were grown under two different Pi regimes, namely 0.05 mM and 0.25 mM Pi (P_0.05_ and P_0.25_ plants). The accumulation of precursor and mature miR827 transcripts substantially decreased when increasing Pi supply, while *NLA* expression increased, thus confirming that the plants respond to Pi treatment (**Fig. S5**).

When growing *miR827prom::GUS* lines plants under 0.05 mM Pi, GUS activity was observed in cotyledons, leaves and roots (**Figure 4**, mock). However, in *MIR827prom::GUS* plants grown under 2 mM Pi, GUS activity was detected only at the distal part of cotyledons and leaves, but not in roots (**Fig. S6A**). To note, treatment with fungal elicitors caused a substantial increase in the activity of the *MIR827* promoter in all Arabidopsis tissues (**Fig. 4**, elicitors), which is in agreement with results obtained by expression analysis (**Fig. 3B**). Furthermore, a remarkable increase in *MIR827* promoter activity occurred in SA-treated *MIR827prom::GUS* plants (**Fig. 4**, SA). Neither Pi treatment (P_0.05_ and P_2_ conditions) (**Fig. S6B)**, nor elicitor or SA-treatment (**Fig. S6C**) had an effect on GUS activity in *35Sprom::GUS* plants.

**Fig. 4.**
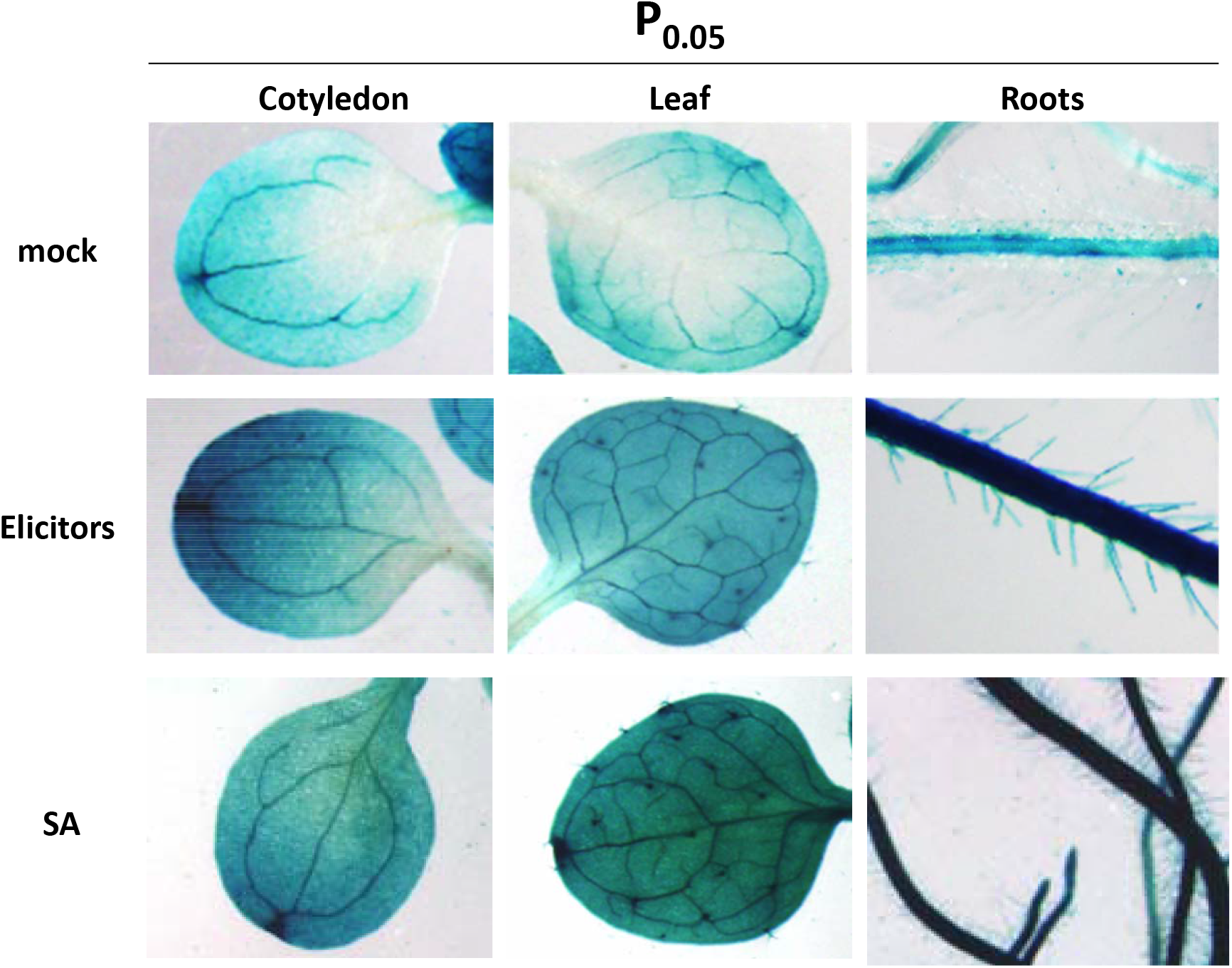
Histochemical analysis of GUS activity in tissues of *miR827prom::GUS* plants that have been grown under 0.05 mM Pi supply. Plants were mock-inoculated or treated with *P. cucumerina* elicitors (300 µg/ml elicitors; 60 min of treatment), or SA (0.1 mM SA; 12 h of treatment). Representative images are shown. Two independent lines were assayed which gave similar results. The analysis of GUS activity in *miR827prom::GUS* plants that have been grown under 0.25mM Pi supply is presented in **Fig. S6**.

The elicitor- and SA-responsiveness that occurs in *MIR827prom::GUS* plants is consistent with the presence of sequence motifs (*cis*-elements) that are known to be involved in pathogen-regulated genes in the *MIR827* promoter (**Fig. S7; Table S1**). In particular, multiple W box TGAC core elements that mediate the transcriptional activation of pathogen- and elicitor-regulated genes, as well as SA-responsive *cis*-elements, were identified in the *MIR827* promoter region (e.g. WBOXATNPR1, TTGAC; ELRECOREPCRP1, TTGACC) (**Fig. S3; Table S1**). As expected, the P1BS element present in phosphate starvation responsive genes was also identified in the *MIR827* promoter (**Fig. S7; Table S1**). Together, these observations point to a transcriptional regulation of *MIR827* expression in response to treatment with fungal elicitors or SA treatment. Transcriptional activation of *MIR827* might well be responsible of down-regulation of *NLA* expression in response to elicitor treatment. However, alterations in *NLA* expression in response to SA treatment remains to be determined.

### Accumulation of camalexin in *nla* mutant plants

Arabidopsis efficiently synthesizes the antifungal phytoalexin camalexin for which an antifungal activity against several pathogens has been reported, including *P. cucumerina* (Ferrari et al., 2003 (Barrlen *et al*., 2007; Sanchez-Vallet *et al*., 2010). To further understand the mechanisms involved in resistance to pathogen infection in nla plants, we examined whether Pi accumulation in Arabidopsis plants leads to alterations in the level of camalexin.

Camalexin originates from tryptophan, and its biosynthesis involves the activity of cytochrome P450 enzymes (**Fig. 5A**). They are: *CYP79B2, CYP79B3, CYP71A13* and *CYP71B15/PAD3*. In this study, we examined the expression of genes involved in the camalexin biosynthetic pathway in *nla* and wild type plants that have that have been inoculated with *P. cucumerina* spores, or mock inoculated. As it is shown in **Fig. 5B**, upon pathogen challenge, all camalexin biosynthesis genes were expressed at a higher level in *nla* plants compared with wild-type plants. Consistent with the observed super-induction of camalexin biosynthesis genes in the *P. cucumerina*-infected *nla* plants, these plants had a higher content of camalexin than wild-type plants (**Fig. 5C**). Finally, an increase in Pi supply in wild type plants is accompanied by a higher level of camalexin, thus, establishing a link between Pi level and camalexin production (**Fig. 5D**). Thus, Pi accumulation in *nla* plants might be responsible of a the observed increase in camalexin accumulation in this plants. This, in turn, would lead to resistance to infection by *P. cucumerina* in Arabidopsis plants.

**Fig. 5.**
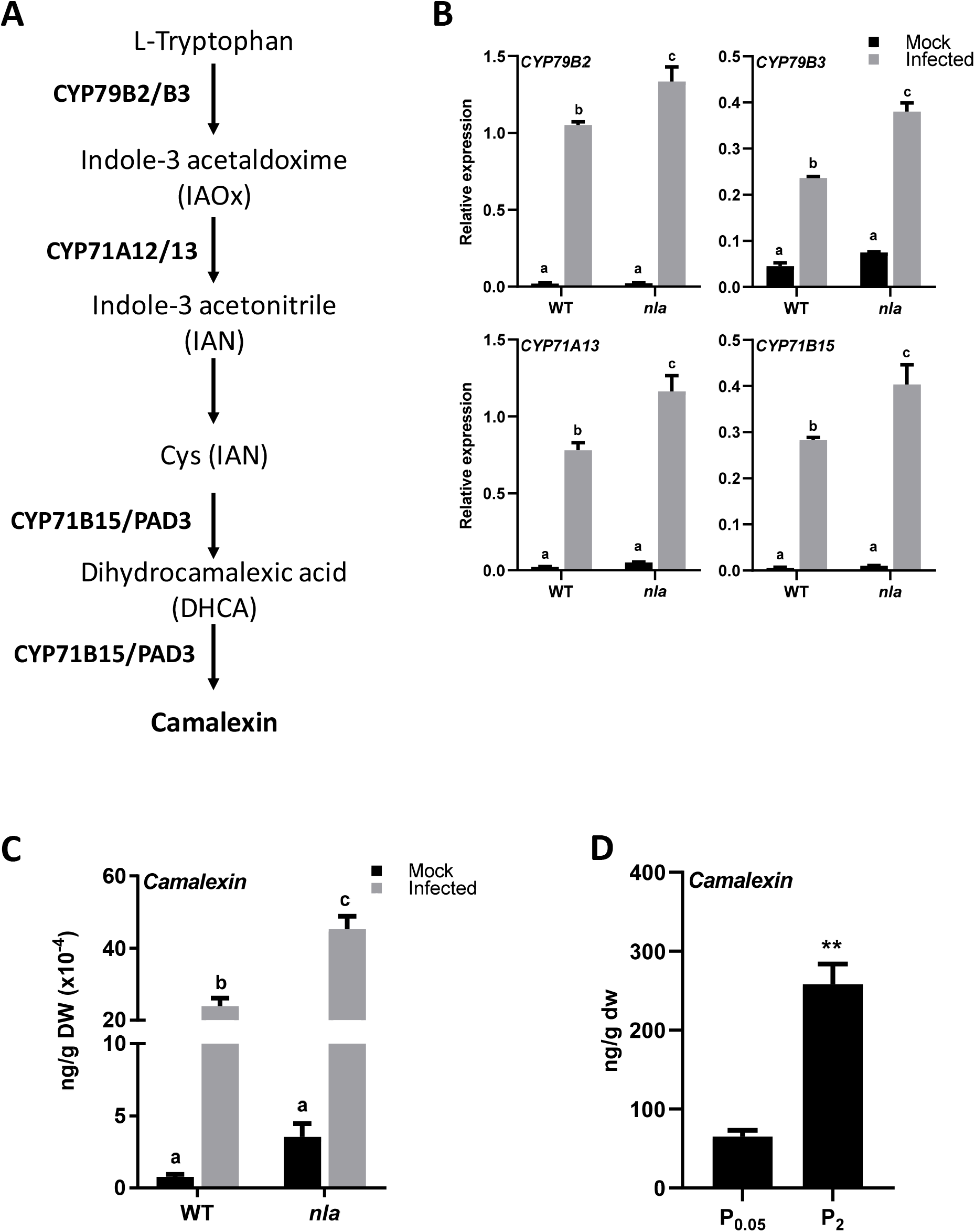
Camalexin accumulation wild type and nla plants and in response to Pi treatment. **A**. Camalexin biosynthesis pathway. **B**. Transcript levels of CYP79B2, CYP79B3, CYP71A13 and CYP71B15 in mock-inoculated and *P. cucumerina*-inoculated wild-type and *nla* plants. Six biological replicates (three plants per replicate were examined). Bars represent mean ± *SEM*. Letters indicate statistically significant differences (ANOVA, HSD Tukey’s test, P ≤ 0.05). **C**. Accumulation of camalexin in wild type and *nla* plants under infection and non infection conditions. **D**. Camalexin accumulation in Pi-treated Arabidopsis plants. Bars represent mean ± *SEM* (six biological replicates and three plants per replicate).

### The Arabidopsis mutant *nla* accumulates SA and JA

In previous studies, it was reported that resistance to *B. cinerea* in Arabidopsis involves camalexin and depends on JA and SA signaling (Ferrari *et al*., 2003). A modulation of camalexin formation by components of the JA signaling pathway was also described (Rowe *et al*., 2010; De Geyter *et al*., 2012). Accordingly, in this work we examined the expression of SA- and JA-regulated defense genes in *nla* plants. In the absence of pathogen infection, the SA markers *PR1* and *NPR1* showed a higher expression in *nla* plants than wild type plants, their expression being further induced upon *P. cucumerina* infection in both genotypes (**Fig. 6A**). Similar levels of pathogen-inducible *PR1* and *NPR1* expression were, however, observed in wild-type and *nla* plants (**Fig. 6A**).

**Fig. 6.**
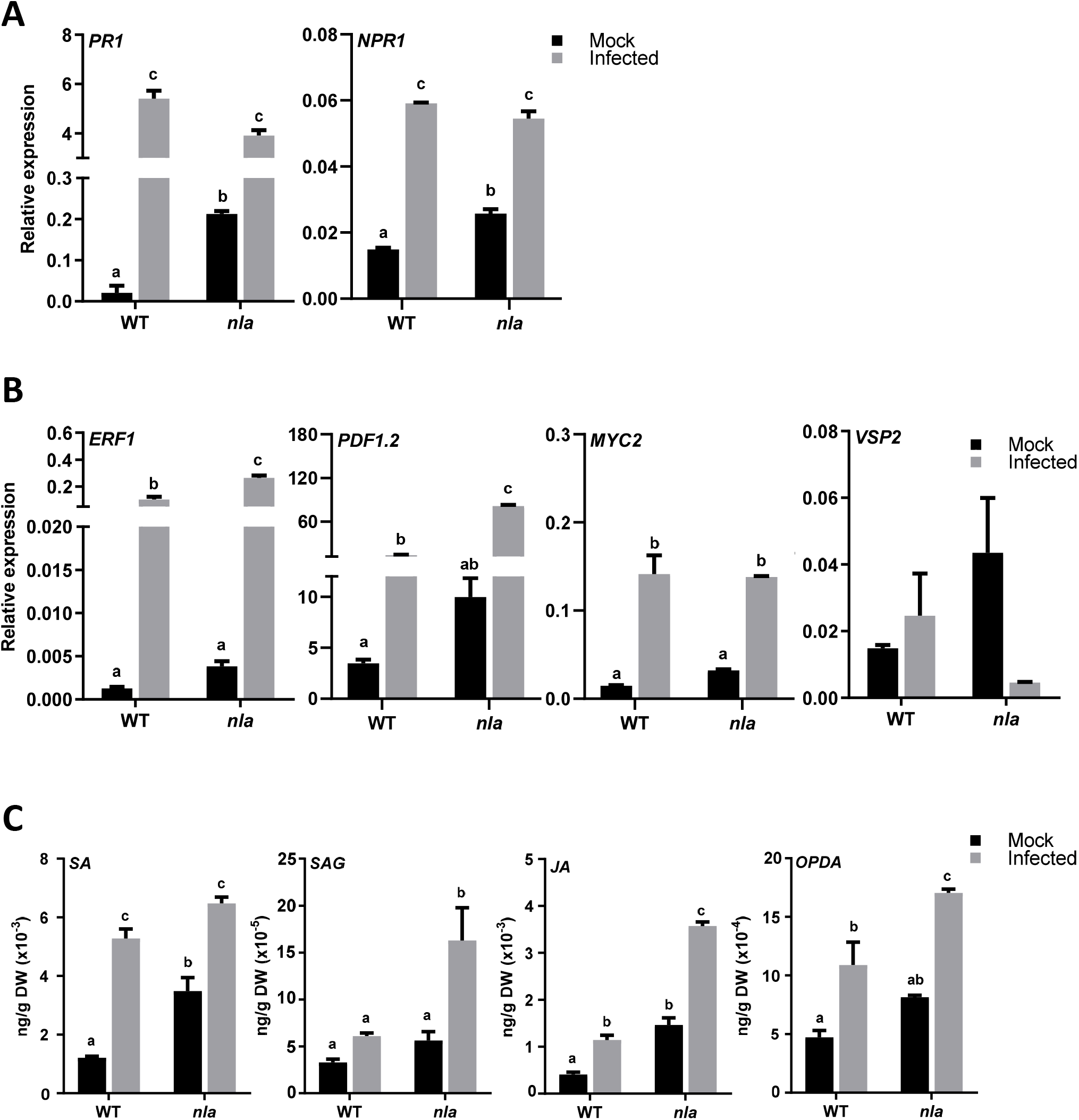
Expression of defense marker genes of the SA and JA signalling pathways and accumulation of SA and JA in wild-type and *nla* plants. Transcript levels in A and B were determined by RT-qPCR in mock-inoculated and *P. cucumerina*-inoculated plants at 48hpi (black and grey bars, respectively). Letters indicate statistically significant differences (ANOVA, HSD Tukey’s test, P ≤ 0.05). **A**. Genes in the SA signaling pathway (*PR1, NPR1*). **B**. Genes in the ERF1/PDF1.2 and MYC2/VSP2 branch in the JA signaling pathway. Bars represent mean ± *SEM* (three biological replicates and three plants per replicate). **C**. Levels of SA, SAG, JA and OPDA. Bars represent mean ± *SEM* (six biological replicates and three plants per replicate).

Next, we examined the expression of genes in the two branches of the JA signaling pathway, the MYC2 branch and ERF branch, which are regulated by AtMYC2 (basic hélix-loop-helix-leucine zipper transcription factor) and AtERF1 (belonging to the APETALA/ERF transcription factor family), respectively (Lorenzo *et al*., 2004). The marker gene commonly used for the ERF branch is *PDF1*.*2* (*Plant Defensin 1*.*2*), whereas the induction of *VSP2* (*Vegetative Storage Protein 2*) serves as marker of the MYC branch (Lorenzo *et al*., 2004; Pieterse *et al*., 2012; Wasternack and Hause, 2013; Zhang *et al*., 2017). Under non-infection conditions, there was a tendency for a higher expression of *ERF1* and *PDF1*.*2* in *nla* plants compared with wild type plants. Equally, a slightly higher *MYC2* and *VSP2* expression occurred in *nla* plants compared with wild-type plants under non infection conditions. *P. cucumerina* infection further induced *ERF1* and *PDF1*.*2*, these genes reaching higher expression level in *nla* plants than in wild type plants (**Fig. 6B**). Regarding the MYC2/VSP2 branch of the JA pathway, *P. cucumerina* infection was found to activate MYC2 expression to a similar level in both genotypes (**Fig. 6B**). In contrast, an opposite response to pathogen infection occurred in *VSP2* expression between wild type and *nla* plants, it expression being up regulated in wild type plants but down-regulated in *nla* plants (**Fig. 6B**).

Then, we measured levels of SA (and the SA glucoside SAG) and JA (and its biosynthetic precursor OPDA) in wild-type and *nla* plants that have been inoculated with *P. cucumerina* or mock-inoculated. In the absence of pathogen infection, *nla* plants accumulated higher levels of SA and JA compared to wild-type plants (**Fig. 6C**). Whereas the pathogen-induced SA level was similar in both genotypes, *P. cucumerina* infection results in increased levels of JA in *nla* plants compared with wild-type plants, (**Fig. 6C**). As for SAG, the storage form of SA, a higher accumulation was observed in *P. cucumerina*-infected *nla* plants relative to the P. *cucumerina*-infected wild type plants pointing to a tight control of SA content in these plants (**Fig. 6C**). The *nla* plants also showed a higher level of OPDA during pathogen infection. As the JA content was significantly increased in *nla* plants under non infection and infection conditions, it is tempting to hypothesize that, in addition to camalexin accumulation, resistance to *P. cucumerina* in *nla* plants also involves JA-dependent defense responses. During pathogen infection, an opposite regulation of the two branches of the JA signaling pathway occur in *nla* plants (e.g up-regulation of *PDF1*.*2* and down-regulation of *VSP2*).

## Discussion

Plants are subjected to a plethora of environmental cues to which they must respond in an efficient and coordinated manner. Upon pathogen infection, plants deploy effective mechanisms to arrest pathogen attack. However, resistance to pathogens can be attenuated or strengthened with co-occurrence of additional stress factors, such as nutrient stress, in particular Pi stress. Although evidence support that Pi deficiency influences the expression of Arabidopsis immune responses, how pathogen-induced signaling pathways converge with signaling pathways triggered by Pi excess still remain obscure.

In this work, we show that an increase in Pi content caused either by loss-of-function of *NLA* or *MIR827* overexpression positively regulates immune responses and confers resistance to infection by fungal pathogens in Arabidopsis. So far, functioning of the miR827/*NLA* pair has been typically associated to the PSR, *NLA* being also involved in adaptation to nitrate limitation in Arabidopsis (Peng *et al*., 2007). Results here presented support that *NLA* also plays a role in Arabidopsis immunity.

Several lines of evidence support that the NLA contributes to disease resistance in Arabidopsis by negatively regulating immune responses. First, loss-of-function of *NLA*, as well as miR827 overexpression, confers resistance to infection. Second, resistance to pathogen infection in *nla* plants is associated with a stronger accumulation of callose. Similarly, Pi treatment in Arabidopsis plants was found to foster callose accumulation during infection with the fungal pathogen *P. cucumerina*. The relevance of callose deposition in resistance to pathogen infection in Arabidopsis, including *P. cucumerina* infection, is well established (Luna *et al*., 2011; Pastor-Fernández *et al*., 2019; Lee *et al*., 2020).Third, *nla* plants showed an increase SA and JA content under non-infection, and accumulated higher levels of JA under infection conditions. To note, an opposite regulation of genes in the two branches of the JA signaling pathway occurs in *nla* and wild-type plants in response to *P. cucumerina* infection. Whereas pathogen infection induces *PDF1*.*2* (marker of the ERF1 branch), *VSP2* (maker of the MYC2 branch) is down regulated by infection in *nla* plants. Fourth, both fungal infection and treatment with fungal elicitors is accompanied by up-regulation of *MIR827* expression and down-regulation of *NLA*. Our results showed that *MIR827* is transcriptionally activated not only by treatment with fungal elicitors but also by treatment with SA, which is consistent with the presence of elicitor- and SA-responsive *cis* elements in the *MIR827* promoter. Several PSI genes have been reported to contain SA-inducible elements in their promoters (Baek et al 2017).

As a further confirmation of the implication of *NLA* in the control of defense responses, the *nla* plants accumulated higher levels of camalexin in their leaves (non-infection and infection conditions) which could be also inferred from the observed superactivation of camalexin biosynthesis genes in these plants. Camalexin plays a key role in Arabidopsis defense against *P. cucumerina*, and the *in vitro* antifungal activity of camalexin against *P. cucumerina* has been demonstrated (Sanchez-Vallet *et al*., 2010). Also, *cyp79B2 cyp79B3* mutants were found to be more susceptible to *P. cucumerina* infection than wild-type plants (Sanchez-Vallet *et al*., 2010). Superinduction of camalexin biosynthesis genes during pathogen infection, and subsequent accumulation of camalexin, might explain the phenotype of disease resistance observed in *nla* mutant plants.

Overall, results here presented on resistance to infection by fungal pathogens in *nla* mutant plants, suggest that *NLA* functions as a negative regulator in Arabidopsis immunity further supporting cross-talk between phosphate signaling, hormone signaling, and disease resistance. JA signaling appears to be important for resistance against P. cucumerina in *nla* plants. In other studies, however, inactivation of *NLA* expression was found to increase susceptibility to the cyst nematode *Heterodera schachtii* (Hewezi *et al*., 2016). Regarding the mechanisms by which loss-of-function of *NLA*, and subsequent accumulation of Pi, might confer resistance to infection by fungal pathogens, several possibilities, not mutually exclusive, can be considered. On the plant side, increased supply of Pi might improve the physiological and metabolic status of the plant which would allow the plant to mount an effective immune response during pathogen infection. Another possibility is that Pi accumulation might have a toxic effect on fungal growth. At present, however, it is not known whether Pi levels might affect *P. cucumerina* (or *C. higginsianum*) growth. The possibility that Pi accumulation might affect disease resistance depending on the type of pathogen and/or pathogen lifestyle in Arabidopsis should be also considered. The observation that *nla* plants exhibit resistance to fungal pathogens with a necrotrophic (*P. cucumerina*) or hemibiotrophic (*C. higginsianum*) lifestyle suggest that *nla*-mediated disease resistance, likely, is not dependent on the pathogen lifestyle. However, contrary to what is observed in Arabidopsis, Pi accumulation in rice was found to increase disease severity caused by the blast fungus *M. oryzae* (Campos-Soriano et al., 2020). Therefore, there is not a general model to predict the role of Pi in a given plant-pathogen interaction. Understanding how Pi signaling pathway and immune signaling interact each other in different plant/pathogen interactions is then of paramount importance in designing strategies to improve disease resistance in plants.

## Conclusions

Taken together, this study suggest that *NLA* functions as a negative regulator of immunity in Arabidopsis. We show that *nla* plants accumulating Pi exhibit stronger immune responses during pathogen infection, such as higher callose deposition and stronger expression of *PR* genes. Disease resistance in *nla* plants might be attributable to the accumulation of the phytoalexin camalexin and increased levels of the defense-related hormones SA (non-infection conditions) and JA (non-infection and infection conditions). This work offered a framework for understanding the impact of Pi accumulation in disease resistance and laid a foundation for future studies to understand interconnected regulations between Pi accumulation and disease resistance.

## Material and Methods

### Plant material infection assays and elicitor treatment

*Arabidopsis thaliana* (ecotype Col 0) were grown under a 12h light/12h dark photoperiod and 60% humidity a temperature of 22ºC ± 2ºC. The *nla* mutant and miR827 OE plants used in this work were previously described (Peng *et al*., 2007, Kant *et al*., 2011, and Lin *et al*., 2013, for *nla*; and Kant et al., 2011, for miR827 OE). For disease resistance assays and elicitor treatment experiments, Arabidopsis plants were grown in a mixture of soil:perlite:vermiculite (2:1:1) and modified Hoagland half strength medium, under neutral photoperiod (12h light / 12h dark), 60% of humidity and a temperature of 22ºC ± 2ºC for three weeks. The fungus *P. cucumerina* was grown on PDA (Potato Dextrose Agar) plates with chloramphenicol (34 µg/ml). *Colletotrichum higginsianum* was grown on Oatmeal agar plates in darkness. Fungal spores were collected by adding sterile water to the surface of the mycelium, and adjusted to the desired final concentration using a Bürker counting chamber. Plants were spray-inoculated with a spore suspension of *P. cucumerina* (5 × 10^5^ spores/ml), or mock-inoculated. *C. higginsianum* was locally-inoculated with a spore suspension at 4 × 10^6^ spores/ml (10 µl/leaf and 5 leaves/plant). Fungal-inoculated and mock-inoculated plants were maintained under high humidity and disease symptom development was followed with time. Lesion area was measured with software ImageJ (National Institute of Health, Bethesda, MD, USA; https://imagej.nih.gov/ij/). Three independent experiments were performed with at least 12 plants per genotype in each experiment. Statistically significant differences were determined by t-test. For *in vitro* experiments, two-week old Arabidopsis plants were spray-inoculated with *P*.*cucumerina* (4 × 10^6^ spores/ml). Fungal biomass was quantified by real-time PCR using specific primers for the corresponding fungus and the Arabidopsis *UBIQUITIN21* (At5g25760) gene as the internal control (Soto-Suárez *et al*., 2017). PCR primers are listed in Table S1. Elicitor treatment was performed by spraying plants with an elicitor extract obtained from *P. cucumerina* (300 μg ml^−1^) as described (Casacuberta *et al*., 1992).

For Pi treatment experiments, plants were grown *in vitro* on meshes placed on agar plates with modified Hoagland half strength medium containing 0.25 mM KH_2_PO_4_ for one week. Seedlings were then transferred to fresh agar-medium at the desired concentration of Pi (0.05, 0.25, or 2 mM Pi). The plants were allowed to continue growing for one more week under each Pi regime. The *in vitro*-grown plants were then inoculated with a spore suspension of *P. cucumerina*.

### Plant tissue staining

For trypan blue staining, leaves were fixed by vacuum infiltration for 1h in ethanol:formaldehyde:acetic acid (80:3.5:5 v/v), stained with lactophenol blue solution for one hour, and then washed with chloral hydrate for 15 minutes. Leaves were placed on glass slides with glycerol and observed using a Leika DM6 microscope under bright field.

Aniline blue staining was used to determine callose deposition. For this, chlorophyll was removed, with 70% ethanol, from leaves that were then incubated in 70 mM phosphate buffer (pH 9.0) supplemented with aniline blue (0.01% wt/vol) with vacuum for 30 min. Samples were maintained in dark conditions for 2 h. Leaves were observed with Leika DM6 microscope under UV illumination. Callose deposition was quantified by determining the relative number of callose-corresponding pixels on digital photographs relative to the total number of pixels using ImageJ software (Luna *et al*., 2011).

For H_2_DCFDA staining, the Arabidopsis leaves were placed on a solution of H_2_DCFDA (at a concentration of 10µM), vacuum infiltrated during 5 minutes, and then maintained in darkness for 10 minutes. Two washes with distillated water were performed. Photographs were taken on a Leika DM6 microscope to visualize green fluorescence. DCFDA staining was quantified by determining the relative number of DCFDA-corresponding pixels relative to the total number of pixels using ImageJ software.

### Agrobacterium-mediated transformation of Arabidopsis

To obtain the *MIR827promoter:GUS* construct, the DNA sequence of the *MIR827* promoter region was extracted from the NCBI (http://www.ncbi.nlm.nih.gov). The transcription start site was identified by using the transcription start site identification program for plants (http://linux1.softberry.com/) and *cis* elements were determined using the PLACE database (https://www.dna.affrc.go.jp/PLACE). The DNA sequence covering 1.2 kb upstream of the transcription start site of *MIR827* was amplified by PCR from genomic DNA, and cloned into the pKGWFS7 plant vector. The PCR product was verified by sequencing. The plant expression vector was transferred to the *Agrobacterium tumefaciens* strain GV3101. Arabidopsis (Col-0) plants were transformed using the floral dip method.

### Analysis of GUS activity

Histochemical staining of GUS enzyme activity was performed according to Jefferson *et al*., 1987. Leaves were fixed by vacuum infiltration for 1 h in ethanol : formaldehyde : acetic acid (80 : 3.5 : 5 by vol.), stained with lactophenol blue solution for 4 h and washed with 70% ethanol (5 min). Leaves were placed on glass slides with glycerol and observed using a Leika DM6 microscope. Hormone treatment with salicylic acid (SA) was performed in a SA solution of 100 µM during 12 hours and darkness.

### Measurements of Pi

The Pi content of Arabidopsis plants was determined as previously described (Versaw and Harrison, 2002). For each experiment four biological replicates (of three plants/replicate) were used. Statistical *t* test analysis was used to analyze the data.

### Gene expression analyses

Total RNA was extracted using TRIzol reagent (Invitrogen). First-strand cDNA was synthesized from DNAse-treated total RNA (1 µg) with reverse transcriptase and oligo-dT (High Capacity cDNA reverse transcription kit, Applied Biosystems). RT-qPCR was performed in optical 96-well plates using SYBR® green in a Light Cycler 480 (Roche). Primers were designed using Primer-Blast (https://www.ncbi.nlm.nih.gov/tools/primer-blast/). The *β-tubulin2* gene (At5g05620) was used to normalize the transcript level in each sample. Primers used for RT-qPCR and stem-loop RT-qPCR are listed in **Supplementary Table S2**. Accumulation of mature miR827 was determined by stem-loop reverse transcription quantitative PCR (Varkonyi-Gasic *et al*., 2007). At least 3 biological replicates were analyzed per genotype and condition, each replicate consisting of leaves from at least 3 independent plants. Two-way analysis of variance (ANOVA) was used to analyze data.

### Determination of Hormones and Camalexin Levels

The rosettes of three-week-old WT (Col-0) and *nla* plants were analyzed by LC-MS for SA, SAG, JA, OPDA and camalexin content as previously described in Sánchez-Bel *et al*., 2018. Briefly, 30 mg of freeze dried material was extracted with MeOH:H2O (30:70) containing 0.01% of HCOOH containing a mixture of 10 ug. L-1 of the internal standards salicylic acid-d 5 (SA-d5) and dehydrojasmonic acid (Sigma-Aldrich). Following extraction, samples were centrifuged (15.000 rpm, 15 min) and filtered through regenerated cellulose filters. An aliquot of 20 ul was injected into a UPLC (Waters Aquity) interfaced with a Xevo TQ-S Mass Spectrometer (TQS, Waters). Hormones were quantified by contrasting with an external calibrarion curve of pure chemical standards of SA, SAG, JA, OPDA and camalexin. Sample separation was performed with a LC Kinetex C18 analytical column of a 5 μm particle size, 2.1 100 mm (Phenomenex). Cinematographic and TQS conditions were performed as described in Sanchez-Bel *et al*. (2018). At least 6 biological replicates were analyzed per genotype and condition, each replicate consisting of leaves from at least 3 independent plants. The plant material was lyophilized prior analysis. Two-way analysis of variance (ANOVA) followed by HSD (Honestly-Significant-Difference) Tukey’s test was used to analyze data. Camalexin levels were determined as previously described (Castillo et al., 2019).

## Supplementary Figures

**Fig. S1.**
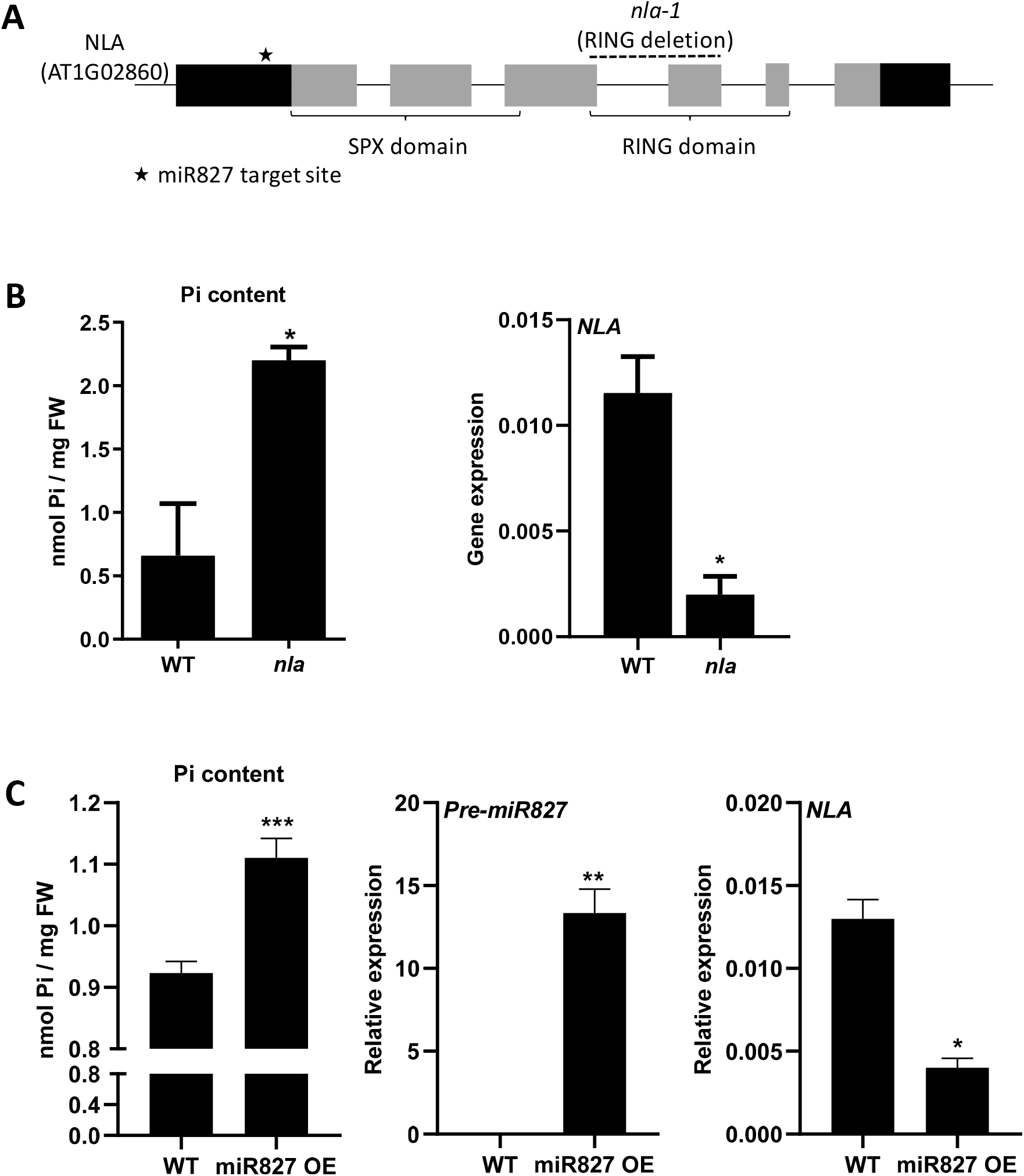
Characterization of *nla* and miR827 OE plants. **A**. Structure of the *NLA* gene. The position of the *nla-1* mutation (deletion) is shown. Black bars, untranslated regions; gray bars, coding regions. **B**. Pi content (left panel) and accumulation of *NLA* transcripts (right panel) in wild-type and *nla* plants. **C**. Accumulation of pre-miR827 and *NLA* transcripts (middle and right panels) and Pi content in miR827 OE plants. The statistical significance was determined by *t* test (\**p* ≤ 0.05, \*\**p* ≤ 0.01 and \*\*\**p* ≤ 0.001).

**Fig. S2.**
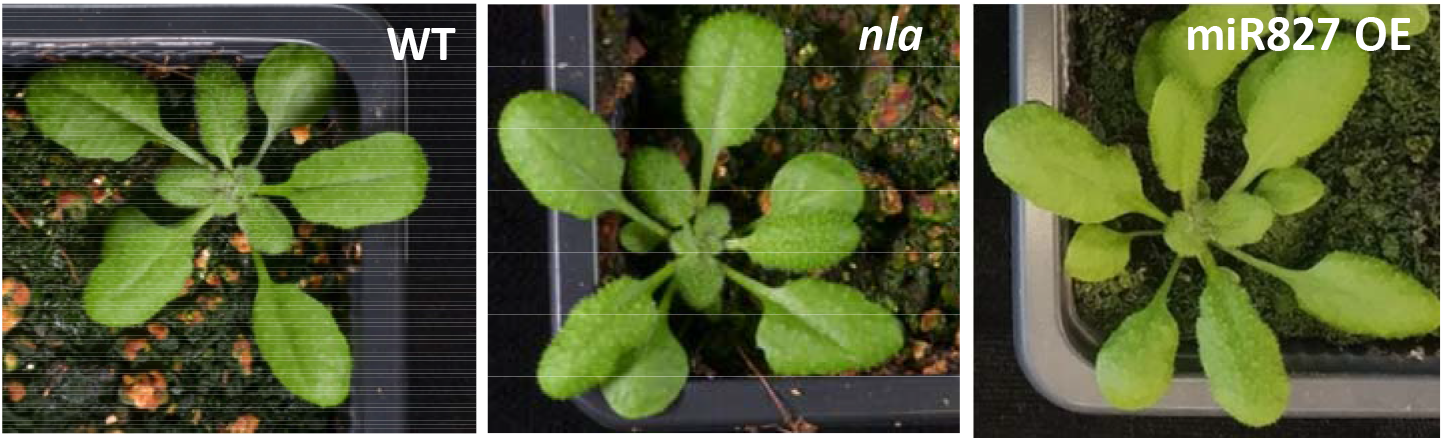
Appearance of 3 week-old wild type, *nla* and miR827 OE plants.

**Fig. S3.**
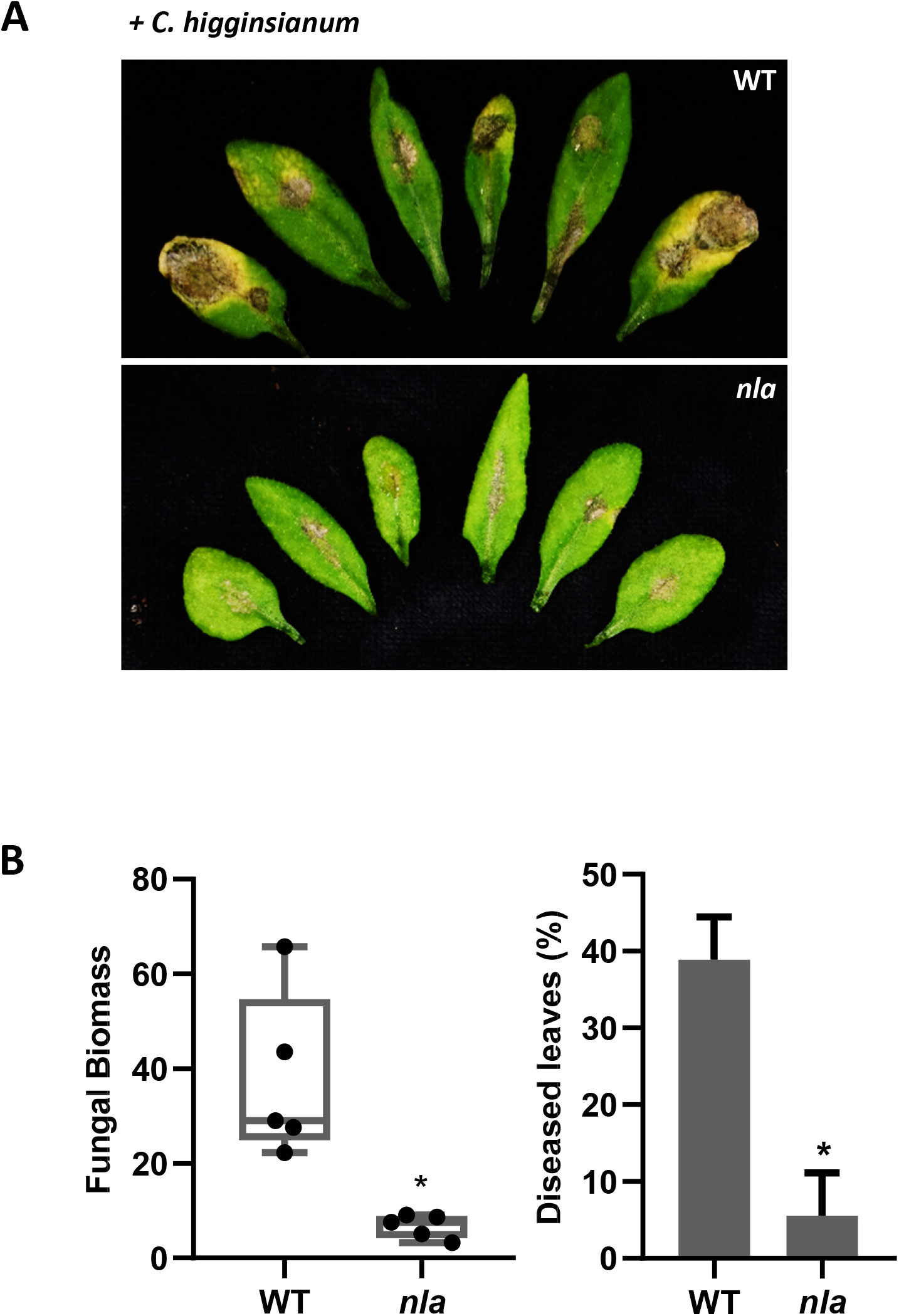
Resistance of *nla* plants to infection by the hemibiotrophic pathogen *C. higginsianum*. Three independent infection experiments were carried out with similar results (at least 24 plants per genotype each experiment). **A**. Phenotype of wild-type and *nla* plants that had been inoculated with *C. higginsianum* spores (4 × 10^6^ spores/ml), or mock inoculated. Pictures were taken at 7 days after inoculation. **B**. Fungal biomass and lesion area was determined at 7 dpi. Quantification of fungal DNA was performed by qPCR using specific primers of *C. higginsianum* (Soto-Suárez *et al*., 2017).

**Fig. S4.**
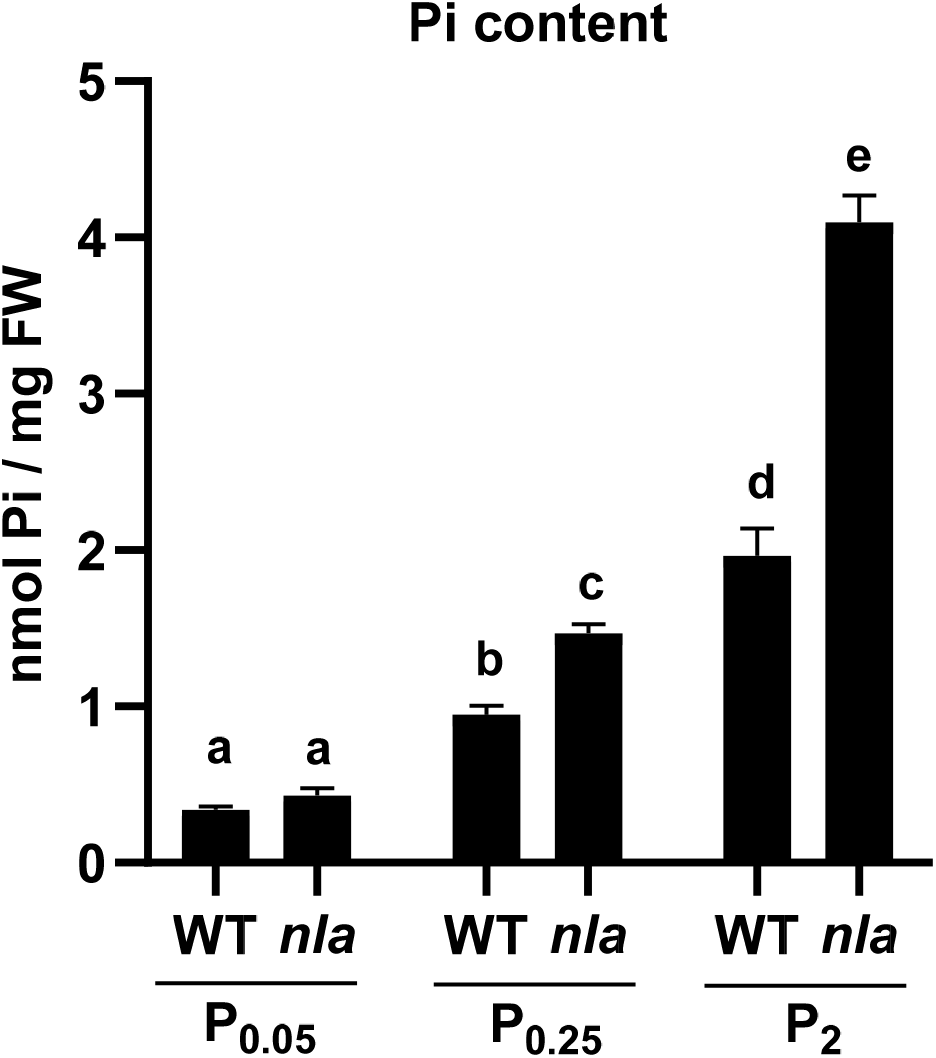
Pi content in wild-type and *nla* plants that have been grown under different Pi supply (0.05 mM, 0.25 and 2 mM Pi).

**Fig. S5.**
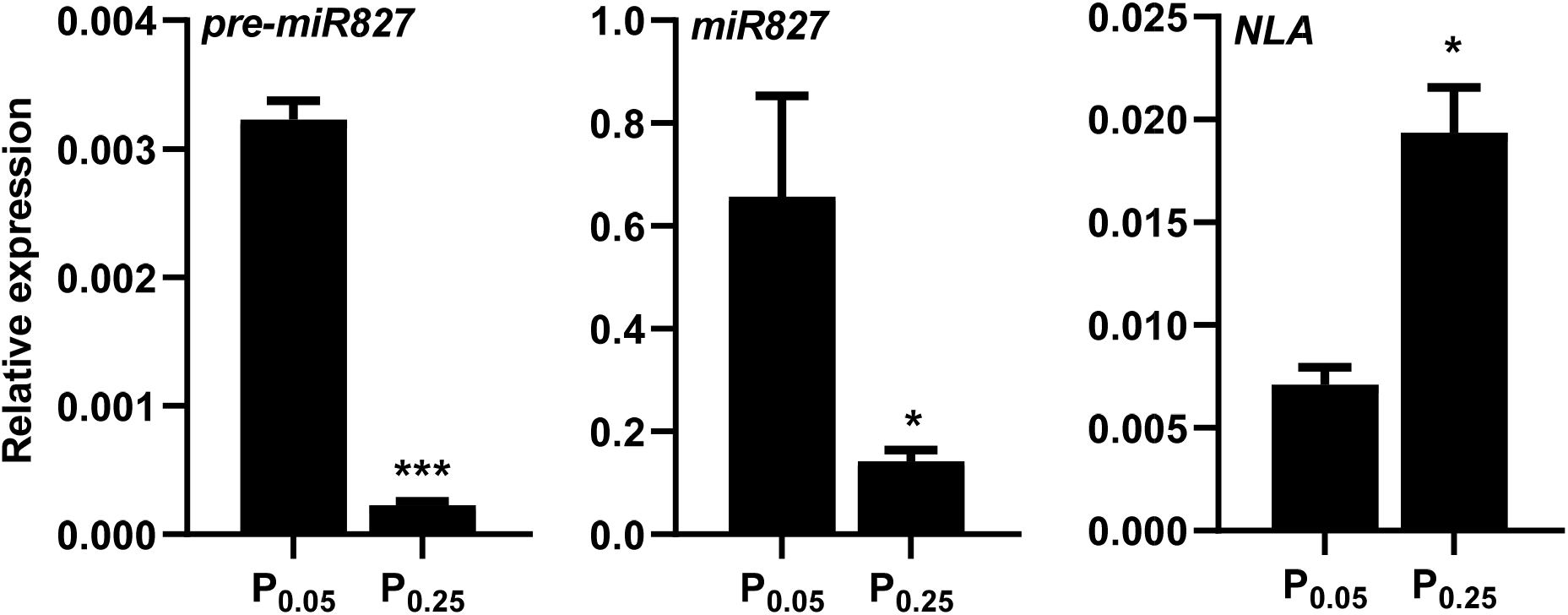
*MIR827* and *NLA* expression in Arabidopsis plants that have been grown under low (P_0.05_) or sufficient (P_0.25_) Pi supply for 7 days. **(A)** Accumulation of pre-miR827, mature miR827 and *NLA* transcripts. **(B)** Pi content in P_0.05_ and P_0.25_ Arabidopsis plants. Bars represent mean ± *SEM* (n=3 and three plants per replicate) (*t* test, \**p* ≤ 0.05, \*\**p* ≤ 0.01 and \*\*\**p* ≤ 0.001).

**Fig. S6.**
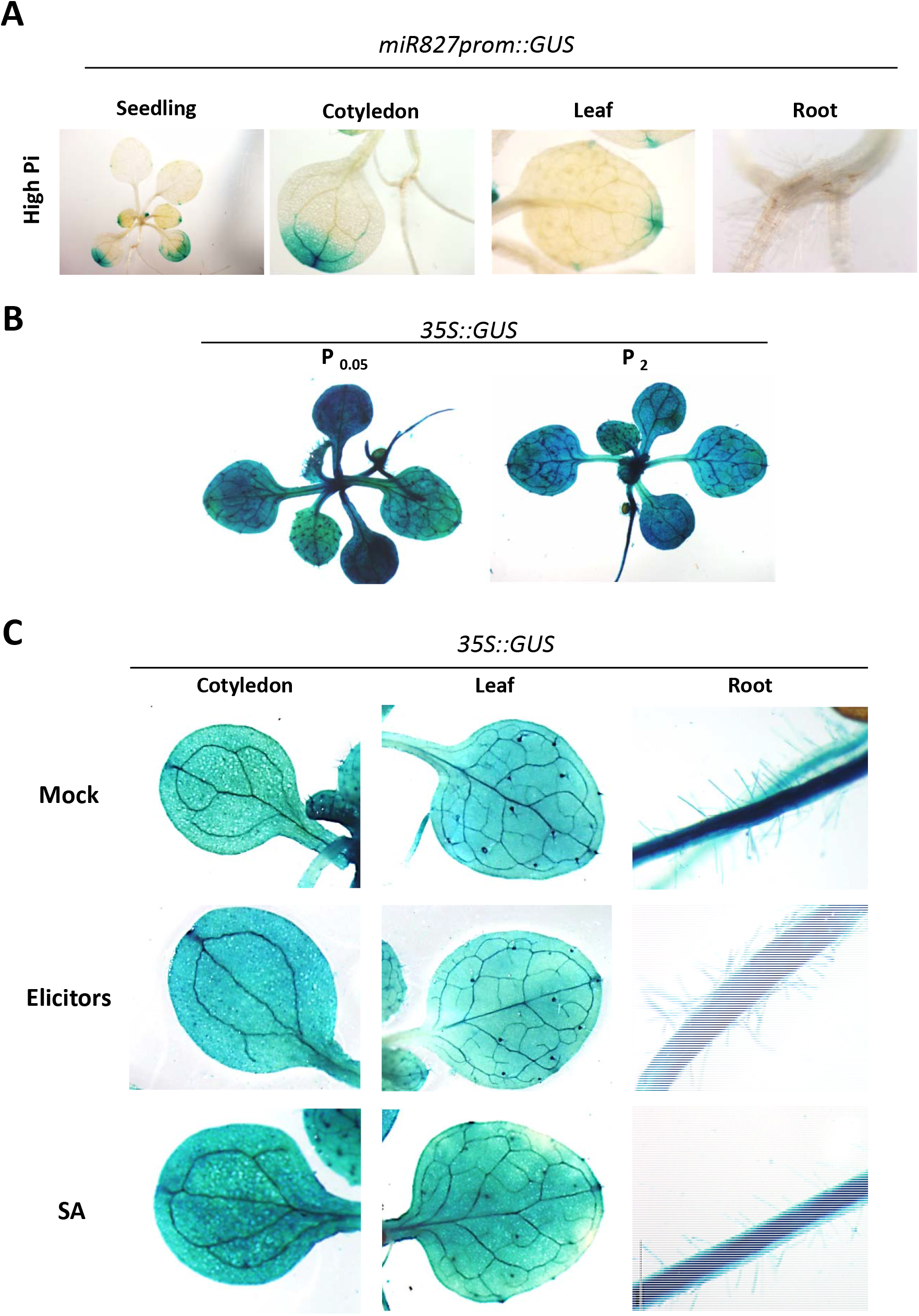
Histochemical analysis of GUS activity of *miR827prom::GUS* and *35S::GUS* Arabidopsis plants. **A**. *miR827prom::GUS* plants grown under high Pi supply (2 mM Pi). **B**. 35S:*:GUS* Arabidopsis plants grown under low or high Pi supply (0.05 mM and 2 mM Pi, respectively). **C**. 35S:*:GUS* plants grown under low Pi supply treated with elicitors prepared from the fungus *P. cucumerina* (0.3 mg/ml; 60 min of treatement) or SA (0.1 mM SA, 12h of treatment). Mock-inoculated plants served as a control.

**Fig. S7.**
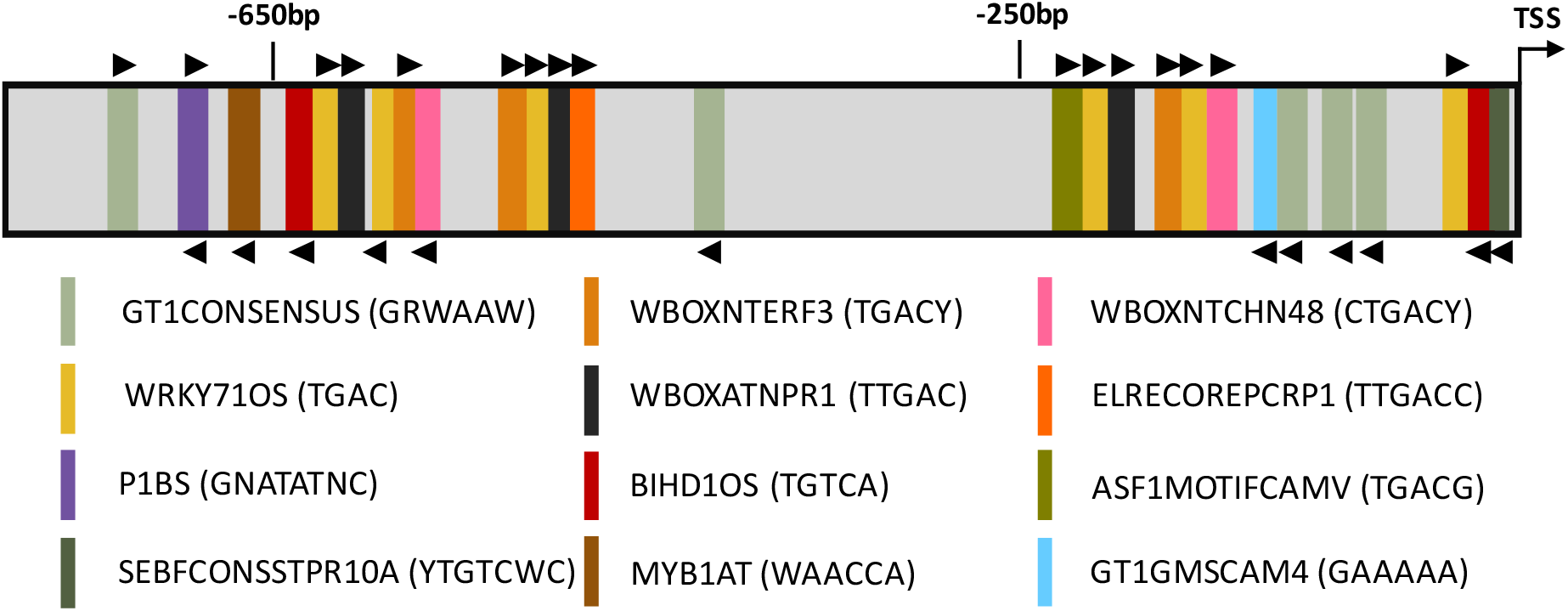
Structural features of the Arabidopsis *MIR827* promoter. The location of known *cis*-acting elements is shown (for details on *cis*-elements, see Supplemental Table S1).

## Supplementary Tables

**Table S1.**
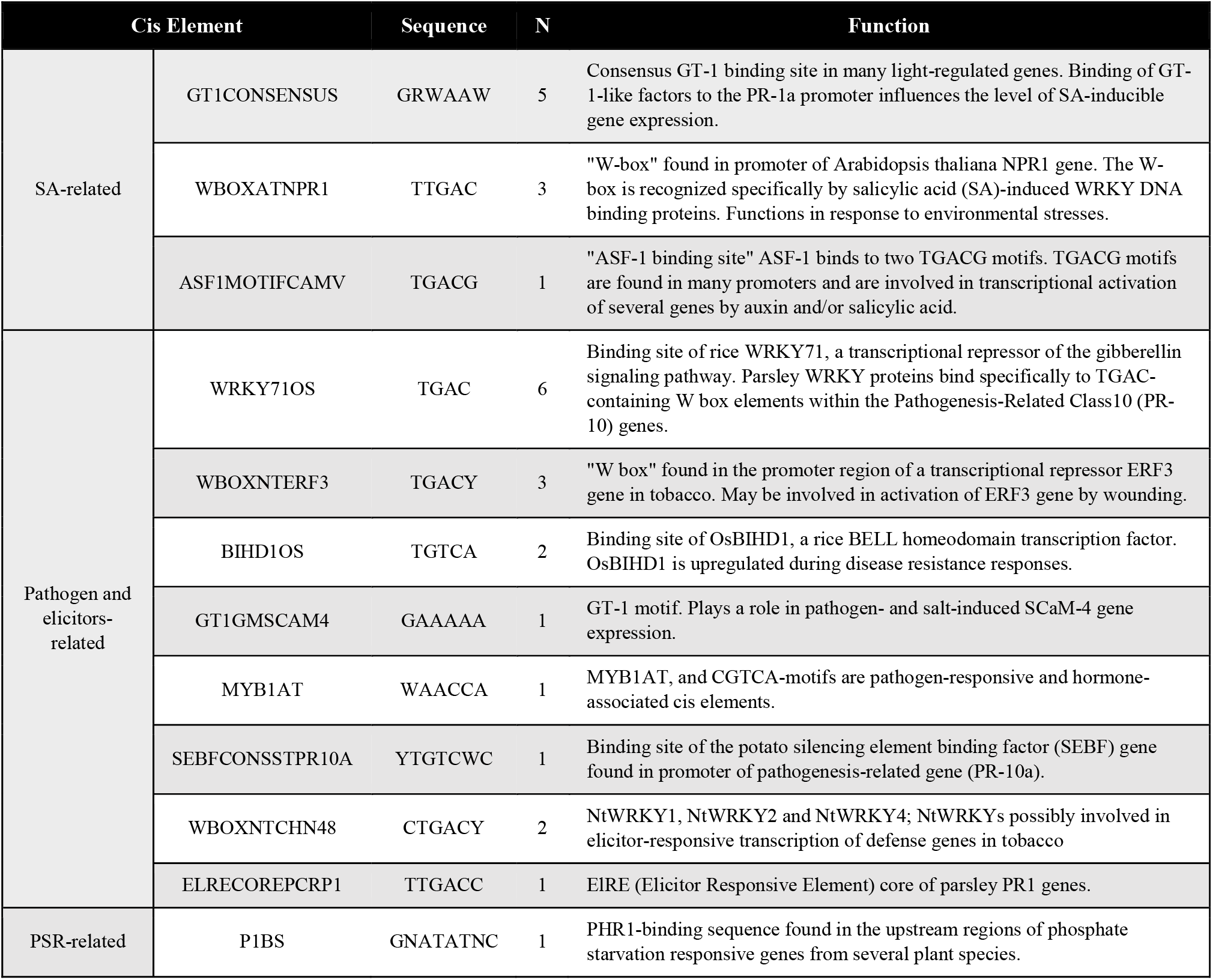
*cis*-elements identified in the MIR827 promoter. The nucleotide sequence, copy number (N) and function of each element are indicated.

**Table S2.**
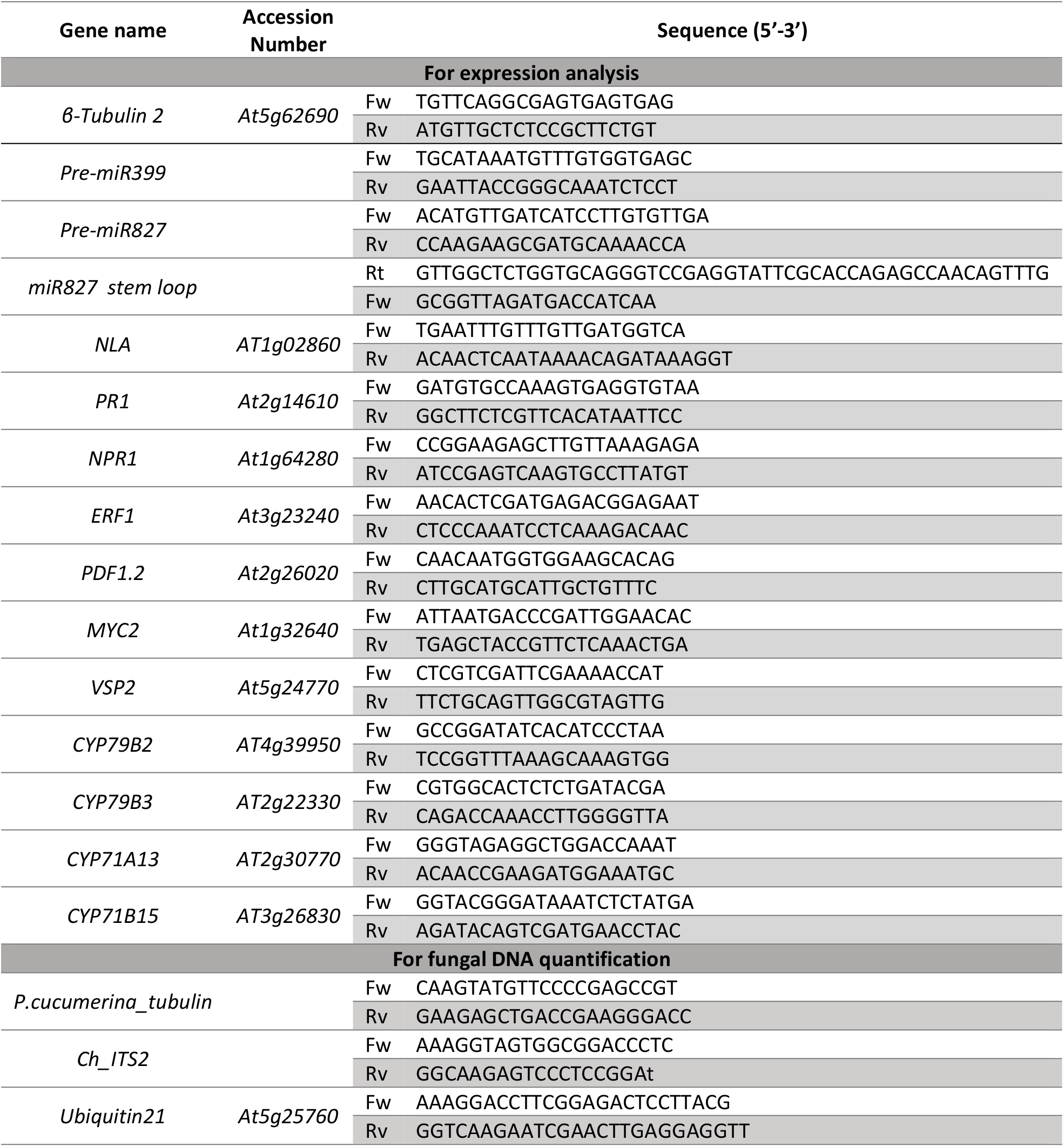
List of oligonucleotides.

## Acknowledgements

This research was supported by FEDER/Ministerio de Ciencia, Innovación y Universidades – Agencia Estatal de Investigación (grant RTI2018-101275-B-I00) to BSS, and Plan de Promoción de la Investigación Universitat Jaume I (UJI-B2019-2) and “Servicios Centrales de Instrumentación Científica (SCIC), Universitat Jaume I to VF. We acknowledge financial support from the Spanish Ministry of Science and Innovation-State Research Agency (AEI), through the “Severo Ochoa Programme for Centres of Excellence in R&D” SEV-2015-0533 and CEX2019-000902-S, and the CERCA Programme / “Generalitat de Catalunya”. M.B.P. was funded by a “La Caixa” scholarship for PhD studies in Spanish universities. B.V-T was a recipient of a Ph. D grant from the Ministerio de Economia, Industria y Competitividad**/**Agencia Estatal de Investigación/Fondo Social Europeo (BES-2016-076289).

## References

Aerts, N., Pereira Mendes, M. and van Wees, S.C.M. (2021) Multiple levels of crosstalk in hormone networks regulating plant defense. Plant J., 105, 489–504.

Ahuja, I., Kissen, R. and Bones, A.M. (2012) Phytoalexins in defense against pathogens. Trends Plant Sci., 17, 73–90.

Baek, D., Chun, H-J., Yun, D-J. and Kim, M-C (2017) Cross-talk between phosphate starvation and other environmental stress signaling pathways in plants Mol. Cells 40, 697–705

Ballini, E., Nguyen, T.T.T. and Morel, J.B. (2013) Diversity and genetics of nitrogen-induced susceptibility to the blast fungus in rice and wheat. Rice, 6, 1–13.

Bari, R., Pant, B.D., Stitt, M. and Scheible, W.R. (2006) PHO2, microRNA399, and PHR1 define a phosphate-signaling pathway in plants. Plant Physiol., 141, 988–999.

Berens, M.L., Berry, H.M., Mine, A., Argueso, C.T. and Tsuda, K. (2017) Evolution of hormone signaling networks in plant defense. Annu. Rev. Phytopathol., 55, annurev–phyto-080516-035544.

Boller, T. and Felix, G. (2009) A renaissance of elicitors: perception of microbe-associated molecular patterns and danger signals by pattern-recognition receptors. Annu. Rev. Plant Biol., 60, 379–406.

Bustos, R., Castrillo, G., Linhares, F., Puga, M.I., Rubio, V., Perez-Perez, J., Solano, R., Leyva, A. and Paz-Ares, J. (2010) A central regulatory system largely controls transcriptional activation and repression responses to phosphate starvation in arabidopsis. PLoS Genet., 6, e1001102.

Campos-Soriano, L., Bundó, M., Bach-Pages, M., Chiang, S.F., Chiou, T.J. and San Segundo, B. (2020) Phosphate excess increases susceptibility to pathogen infection in rice. Mol. Plant Pathol., 21, 555–570.

Casacuberta, J.M., Raventós, D., Puigdoménech, P. and Segundo, B.S. (1992) Expression of the gene encoding the PR-like protein PRms in germinating maize embryos. Mol. Gen. Genet., 234, 97–104.

Castillo, N., Pastor, V., Chávez, A., Arró, M., Boronat, A., Flors, V., Ferrer, A., and Altabella, T. (2019) Inactivation of UDP-Glucose Sterol Glucosyltransferases Enhances Arabidopsis Resistance to Botrytis cinerea. Fr. Pl. Sci. 10:1162

Castrillo, G., Teixeira, P.J.P.L., Paredes, S.H., et al. (2017) Root microbiota drive direct integration of phosphate stress and immunity. Nature, 543, 513–518.

Chan, C., Liao, Y.-Y. and Chiou, T.-J. (2021) The impact of phosphorus on plant immunity. Plant Cell Physiol. 62, 582–589.

Chien, P-S., Chiang, C-P., Leong S-J., Chiou, T-J (2018) Sensing and Signaling of Phosphate Starvation: From Local to Long Distance. Plant Cell Physiol. 59, 1714–1722

Delhaize, E. and Randall, P.J. (1995) Characterization of a phosphate-accumulator mutant of Arabidopsis thaliana. Plant Physiol., 107, 207–213.

Denancé, N., Sánchez-Vallet, A., Goffner, D. and Molina, A. (2013) Disease resistance or growth: the role of plant hormones in balancing immune responses and fitness costs. Front. Plant Sci., 4, 1–12.

Ferrari, S., Plotnikova, J.M., De Lorenzo, G., and Ausubel, F.M. (2003) Arabidopsis local resistance to Botrytis cinerea involves salicylic acid and camalexin and requires EDS4 and PAD2, but not SID2, EDS5 or PAD4. Plant J., 35, 193–205.

Guo, M., Ruan, W., Li, C., et al. (2015) Integrative comparison of the role of the PHOSPHATE RESPONSE1 subfamily in phosphate signaling and homeostasis in rice. Plant Physiol., 168, 1762–1776.

Hewezi, T., Piya, S., Qi, M., Balasubramaniam, M., Rice, J.H. and Baum, T.J. (2016) Arabidopsis miR827 mediates post-transcriptional gene silencing of its ubiquitin E3 ligase target gene in the syncytium of the cyst nematode Heterodera schachtii to enhance susceptibility. Plant J., 88, 179–192.

Hsieh, L.C., Lin, S.I., Shih, A.C.C., Chen, J.W., Lin, W.Y., Tseng, C.Y., Li, W.H. and Chiou, T.J. (2009) Uncovering small RNA-mediated responses to phosphate deficiency in Arabidopsis by deep sequencing. Plant Physiol., 151, 2120–2132.

Jefferson, R.A., Kavanagh, T.A. and Bevan, M.W. (1987) GUS fusions: beta-glucuronidase as a sensitive and versatile gene fusion marker in higher plants. EMBO J., 6, 3901.

Jones, J.D.G. and Dangl, J.L. (2006) The plant immune system. Nature, 444, 323–329.

Kant, S., Peng, M. and Rothstein, Steven J. (2011) Genetic Regulation by NLA and microRNA827 for maintaining nitrate-dependent phosphate homeostasis in Arabidopsis L.-J. Qu, ed. PLoS Genet., 7, e1002021.

Lee, D.H., Lal, N.K., Lin, Z.J.D., Ma, S., Liu, J., Castro, B., Toruño, T., Dinesh-Kumar, S.P. and Coaker, G. (2020) Regulation of reactive oxygen species during plant immunity through phosphorylation and ubiquitination of RBOHD. Nat. Commun., 11, 1–16.

Li, P., Lu, Y.-J., Chen, H. and Day, B. (2020) The lifecycle of the plant immune system. Cri. Rev. PLant. Sci., 39, 72–100.

Lin, W.Y., Huang, T.K. and Chiou, T.J. (2013) NITROGEN LIMITATION ADAPTATION, a target of microRNA827, mediates degradation of plasma membrane-localized phosphate transporters to maintain phosphate homeostasis in Arabidopsis. Plant Cell, 25, 4061–4074.

Lorenzo, O., Chico, J.M., Sánchez-Serrano, J.J. and Solano, R. (2004) JASMONATE-INSENSITIVE1 encodes a MYC transcription factor essential to discriminate between different jasmonate-regulated defense responses in Arabidopsis. Plant Cell, 16, 1938–1950.

Lu, Y.T., Li, M.Y., Cheng, K.T., Tan, C.M., Su, L.W., Lin, W.Y., Shih, H.T., Chiou, T.J. and Yang, J.Y. (2014) Transgenic plants that express the phytoplasma effector SAP11 show altered phosphate starvation and defense responses. Plant Physiol., 164, 1456–1469.

Luna, E., Pastor, V., Robert, J., Flors, V., Mauch-Mani, B. and Ton, J. (2011) Callose deposition: a multifaceted plant defense response. Mol. Plant-Microbe Interact., 24, 183–193.

Nafisi, M., Goregaoker, S, Botanga, C.J., Glawischnig, E., Olsen, C.E., Halkier, B.A., and Glazebrook, J. (2007) Arabidopsis cytochrome P450 monooxygenase 71A13 catalyzes the conversion of indole-3-acetaldoxime in camalexin synthesis. Plant Cell, 19, 2039–2052.

Nilsson, L., Lundmark, M., Jensen, P.E. and Nielsen, T.H. (2012) The Arabidopsis transcription factor PHR1 is essential for adaptation to high light and retaining functional photosynthesis during phosphate starvation. Physiol. Plant., 144, 35–47.

O’Connell, R., Herbert, C., Sreenivasaprasad, S., Khatib, M., Esquerré-Tugayé, M.T. and Dumas, B. (2004) A novel Arabidopsis-Colletotrichum pathosystem for the molecular dissection of plant-fungal interactions. Mol. Plant-Microbe Interact., 17, 272–282.

Pastor, V., Luna, E., Ton, J., Cerezo, M., García-Agustín, P., and Flors, V (2013) Fine tuning of reactive oxygen species homeostasis regulates primed immune responses in Arabidopsis. Mol. Plant-Microbe Interact. 26, 1334–1344.

Pastor-Fernández, J., Pastor, V., Mateu, D., Gamir, J., Sánchez-Bel, P. and Flors, V. (2019) Accumulating evidences of callose priming by indole-3-carboxylic acid in response to Plectospharella cucumerina. Plant Signal. Behav., 14, 1608107.

Peng, M., Hannam, C., Gu, H., Bi, Y.M. and Rothstein, S.J. (2007) A mutation in NLA, which encodes a RING-type ubiquitin ligase, disrupts the adaptability of Arabidopsis to nitrogen limitation. Plant J., 50, 320–337.

Pieterse, C.M.J., Does, D. Van Der, Zamioudis, C., Leon-Reyes, A. and Wees, S.C.M. Van (2012) Hormonal modulation of plant immunity. Annu. Rev. Cell Dev. Biol., 28, 489–521.

Puga, M.I., Rojas-Triana, M., Lorenzo, L. de, Leyva, A., Rubio, V. and Paz-Ares, J. (2017) Novel signals in the regulation of Pi starvation responses in plants: facts and promises. Curr. Opin. Plant Biol., 39, 40–49.

Rowe, H.C., Walley, J.W., Corwin, J., Chan, E.K.-F., Dehesh, K. and Kliebenstein, D.J. (2010) Deficiencies in jasmonate-mediated plant defense reveal quantitative variation in Botrytis cinerea pathogenesis. PLoS Pathog., 6, 1–18.

Rubio, V., Linhares, F., Solano, R., Martín, A.C., Iglesias, J., Leyva, A. and Paz-Ares, J. (2001) A conserved MYB transcription factor involved in phosphate starvation signaling both in vascular plants and in unicellular algae. Genes Dev., 15, 2122–2133.

Saijo, Y. and Loo, E.P. (2020) Plant immunity in signal integration between biotic and abiotic stress responses. New Phytol., 225, 87–104.

Sánchez-Bel, P., Sanmartín, N., Pastor, V., Mateu, D., Cerezo, M., Vidal-Albalat, A., Pastor-Fernández, J., Pozo, M.J. and Flors, V. (2018) Mycorrhizal tomato plants fine tunes the growth-defence balance upon N depleted root environments. Plant Cell Environ., 41, 406–420.

Sánchez-Vallet, A., Ramos, B., Bednarek, P., López, G., Piślewska-Bednarek, M., Schulze-Lefert, P. and Molina, A. (2010) Tryptophan-derived secondary metabolites in Arabidopsis thaliana confer non-host resistance to necrotrophic Plectosphaerella cucumerina fungi. Plant J., 63, 115–127.

Soto-Suárez, M., Baldrich, P., Weigel, D., Rubio-Somoza, I. and San Segundo, B. (2017) The Arabidopsis miR396 mediates pathogen-associated molecular pattern-triggered immune responses against fungal pathogens. Sci. Rep., 7, 1–14.

van Baarlen, P., Woltering, E.J., Staats, M. and van Kan, J.A.L. (2007) Histochemical and genetic analysis of host and non-host interactions of Arabidopsis with three Botrytis species: an important role for cell death control. Mol. Plant Pathol., 8, 41–54.

Varkonyi-Gasic, E., Wu, R., Wood, M., Walton, E.F. and Hellens, R.P. (2007) Protocol: A highly sensitive RT-PCR method for detection and quantification of microRNAs. Plant Methods, 3, 1–12.

Veresoglou, S.D., Barto, E.K., Menexes, G. and Rillig, M.C. (2013) Fertilization affects severity of disease caused by fungal plant pathogens. Plant Pathol., 62, 961–969.

Versaw, W.K. and Harrison, M.J. (2002) A chloroplast phosphate transporter, PHT2;1, influences allocation of phosphate within the plant and phosphate-starvation responses. Plant Cell, 14, 1751–1766.

Wang, Y., Li, X., Fan, B., Zhu, C. and Chen, Z. (2021) Regulation and function of defense-related callose deposition in plants. Int. J. Mol. Sci., 22, 2393.

Wasternack, C. and Hause, B. (2013) Jasmonates: Biosynthesis, perception, signal transduction and action in plant stress response, growth and development. An update to the 2007 review in Annals of Botany. Ann. Bot., 111, 1021–1058.

Wang, Y., Wang, F., Lu, H., Liu, Y. and Mao, C. (2021) Phosphate uptake and transport in plants: An elaborate regulatory system. Plant Cell Physiol. 62, 564–572.

Yaeno, T. and Iba, K. BAH1/NLA, a RING-type ubiquitin E3 ligase, regulates the accumulation of salicylic acid and immune responses to Pseudomonas syringae DC3000 1. Plant Physiol., 148, 1032–1041.

Yang, X.J. and Finnegan, P.M. (2010) Regulation of phosphate starvation responses in higher plants. Ann. Bot., 105, 513–526.

Zhang, L., Zhang, F, Melotto, M., Yao, J. and He, S-Y (2017) Jasmonate signaling and manipulation by pathogens and insects. J. of Exp. Bot. 68, 1371–1385

